# A conserved strategy for inducing appendage regeneration

**DOI:** 10.1101/2020.11.21.392720

**Authors:** Michael J. Abrams, Fayth Tan, Ty Basinger, Martin Heithe, Yutian Li, Misha Raffiee, Patrick Leahy, John O. Dabiri, David A. Gold, Lea Goentoro

## Abstract

Can limb regeneration be induced? Few have pursued this question, and an evolutionarily conserved strategy has yet to emerge. This study reports a strategy for inducing regenerative response in appendages, which works across three species that span the animal phylogeny. In Cnidaria, the frequency of appendage regeneration in the moon jellyfish *Aurelia* was increased by feeding with the amino acid L-leucine and the growth hormone insulin. In insects, the same strategy induced tibia regeneration in adult *Drosophila*. Finally, in mammals, L-leucine and sucrose administration induced digit regeneration in adult mice, including dramatically from mid-phalangeal amputation. The conserved effect of L-leucine and insulin/sugar suggests a key role for energetic parameters in regeneration induction. The simplicity by which nutrient supplementation can induce appendage regeneration provides a testable hypothesis across animals.

## Introduction

In contrast to humans’ poor ability to regenerate, the animal world is filled with seemingly Homeric tales: a creature that regrows when halved or a whole animal growing from a small body piece. Two views have historically prevailed as to why some animals regenerate better than others (Goss, 1992; Polezhaev, 1972; Morgan, 1901). Some biologists, including Charles Darwin and August Weismann, hold that regeneration is an adaptive property of a specific organ. For instance, some lobsters may evolve the ability to regenerate claws because they often lose them in fights and food foraging. Other biologists, including Thomas Morgan, hold that regeneration is not an evolved trait of a particular organ, but inherent in all organisms. Regeneration evolving for a particular organ versus regeneration being organismally inherent is an important distinction, as the latter suggests that the lack of regeneration is not due to the trait never having evolved, but rather due to inactivation – and may therefore be induced. In support of Morgan’s view, studies in past decades have converged on one striking insight: many animal phyla have at least one or more species that regenerate body parts (Sanchez-Alvarado, 2000; Bely and Nyberg, 2010). Further, even in poorly regenerative lineages, many embryonic and larval stages can regenerate. In fish, conserved regeneration-responsive enhancers were recently identified, which are also modified in mice (Wang et al., 2020). These findings begin to build the case that, rather than many instances of convergence, the ability to regenerate is ancestral (Sanchez-Alvarado, 2000; Bely and Nyberg, 2010). Regeneration being ancestral begs the question: is there a conserved mechanism to activate regenerative state?

This study explored how, and whether, limbs can be made to regenerate in animals that do not normally show limb regeneration. In frogs, studies from the early 20th century and few recent ones have induced various degrees of outgrowth in the limb using strategies including repeated trauma, electrical stimulation, local progesterone delivery, progenitor cell implantation, and Wnt activation (Carlson, 2007; Lin et al., 2013; Kawakami et al., 2006). Wnt activation restored limb development in chick embryos (Kawakami et al., 2006), but there are no reports of postnatal regeneration induction. In salamanders, a wound site that normally just heals can be induced to grow a limb by supplying nerve connection and skin graft from the contralateral limb (Endo et al., 2004), or by delivery of Fgf2, 8, and Bmp2 to the wound site followed by retinoic acid (Viera et al., 2019). In mouse digits, a model for exploring limb regeneration in mammals, bone outgrowth or joint-like structure can be induced via local implantation of Bmp2 or 9 (Yu et al., 2019). Thus far, different strategies gain tractions in different species, and a common denominator appears elusive.

However, across animal phylogeny, some physiological features show interesting correlation with regenerative ability (Hariharan et al., 2015; Vivien et al., 2016; Sousounis et al., 2014). First, regeneration tends to decrease with age, with juveniles and larvae more likely to regenerate than adults. For instance, the mammalian heart rapidly loses the ability to regenerate after birth and anurans cease to regenerate limbs upon metamorphosis. Second, animals that continue to grow throughout life tend to also regenerate. For instance, most annelids continue adding body segments and regenerate well, a striking exception of which is leeches that make exactly 32 segments and one of the few annelids that do not regenerate body segments. Consistent with the notion of regeneration as ancestral, indeterminate growth is thought of as the ancestral state (Hariharan et al., 2015). Finally, a broad correlate of regenerative ability across animal phylogeny is thermal regulation. Poikilotherms, which include most invertebrates, fish, reptiles and amphibians, tend to have greater regenerative abilities than homeotherms — birds and mammals are animal lineages with poorest regeneration. These physiological correlates, taken together, are united by the notion of energy expenditure. The transition from juvenile to adult is a period of intense energy usage, continued growth is generally underlined by sustained anabolic processes, and regulating body temperature is energetically expensive compared to allowing for fluctuation. Regeneration itself entails activation of anabolic processes to rebuild lost tissues (Hirose et al., 2014; Naviaux et al., 2009; Malandraki-Miller et al., 2018). These physiological correlates thus raise the notion of a key role of energetics in the evolution of regeneration in animals. Specifically, we wondered whether energy inputs can promote regenerative state. In this study, we demonstrate that nutrient supplementation can induce regenerative response in appendage and limb across three vastly divergent species.

## Results

### Leucine and insulin promote appendage regeneration in the moon jelly *Aurelia*

We reasoned that if there was an ancestral mechanism to promote regeneration, it would more likely be intact in early-branching lineages. In Cnidaria, the ability to regenerate is established in polyps, *e.g.*, hydras and sea anemones. Some cnidarians, notably jellyfish, not only exist as sessile polyps, but also as free-swimming ephyrae and medusae (Figure 1a). In contrast to the polyps’ ability to regenerate, regeneration in ephyrae and medusae appears more restricted (Abrams et al., 2015). We focused on the moon jellyfish *Aurelia coerulea* (formally *A. aurita* sp. 1 strain), specifically on the ephyra, whose eight arms facilitate morphological tracking (Figure 1b). *Aurelia* ephyrae regenerate tips of arms and the distal sensory organ rhopalium, but upon more dramatic amputations such as removing a whole arm or halving the body, rapidly reorganize existing body parts and regain radial symmetry (Figure 1c). Observed across four scyphozoan species, symmetrization occurs rapidly within 1-3 days and robustly across conditions (Abrams et al., 2015). Ephyrae that symmetrized matured into medusae, whereas ephyrae that failed to symmetrize and simply healed the wound grew abnormally.

**Figure 1.**
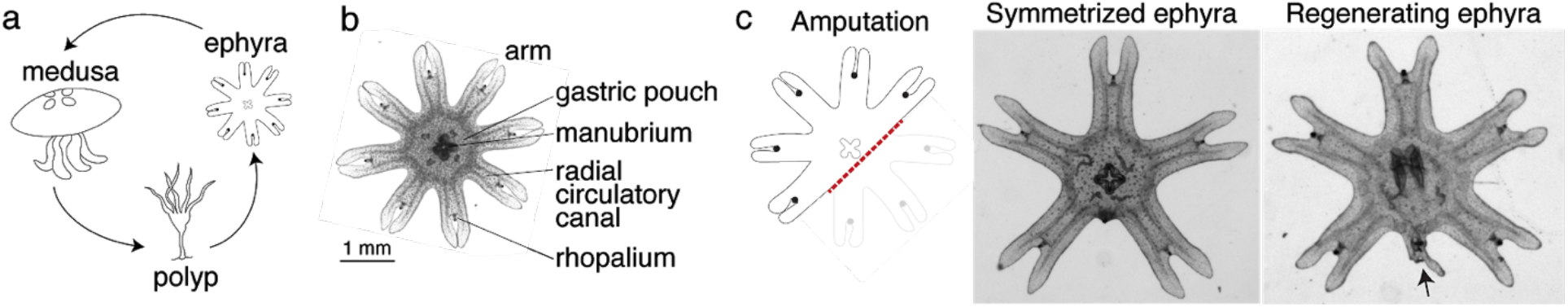
*Aurelia* as a system to identify factors that promote appendage regeneration. **(a)** The moon jellyfish *Aurelia aurita* have a dimorphic life cycle, existing as sessile polyps or free-swimming medusae and ephyrae. Ephyra is the juvenile stage of medusa, a robust stage that can withstand months of starvation. In lab conditions, ephyrae mature into medusae, growing bell tissue and reproductive organs, in 1-2 months. **(b)** Ephyrae have eight arms, which are swimming appendages that contract synchronously to generate axisymmetric fluid flow, which facilitates propulsion and filter feeding. The eight arms are symmetrically positioned around the stomach and the feeding organ manubrium. Extending into each arm is radial muscle (shown in Figure 2) and a circulatory canal that transports nutrients. At the end of each arm is the light- and gravity-sensing organ rhopalium. **(c)** In response to injury, the majority of ephyrae rapidly reorganize existing body parts and regain radial symmetry. However, performing the experiment in the natural habitat, a few ephyrae (2 of 18) regenerated a small arm (arrow).

**Figure 2.**
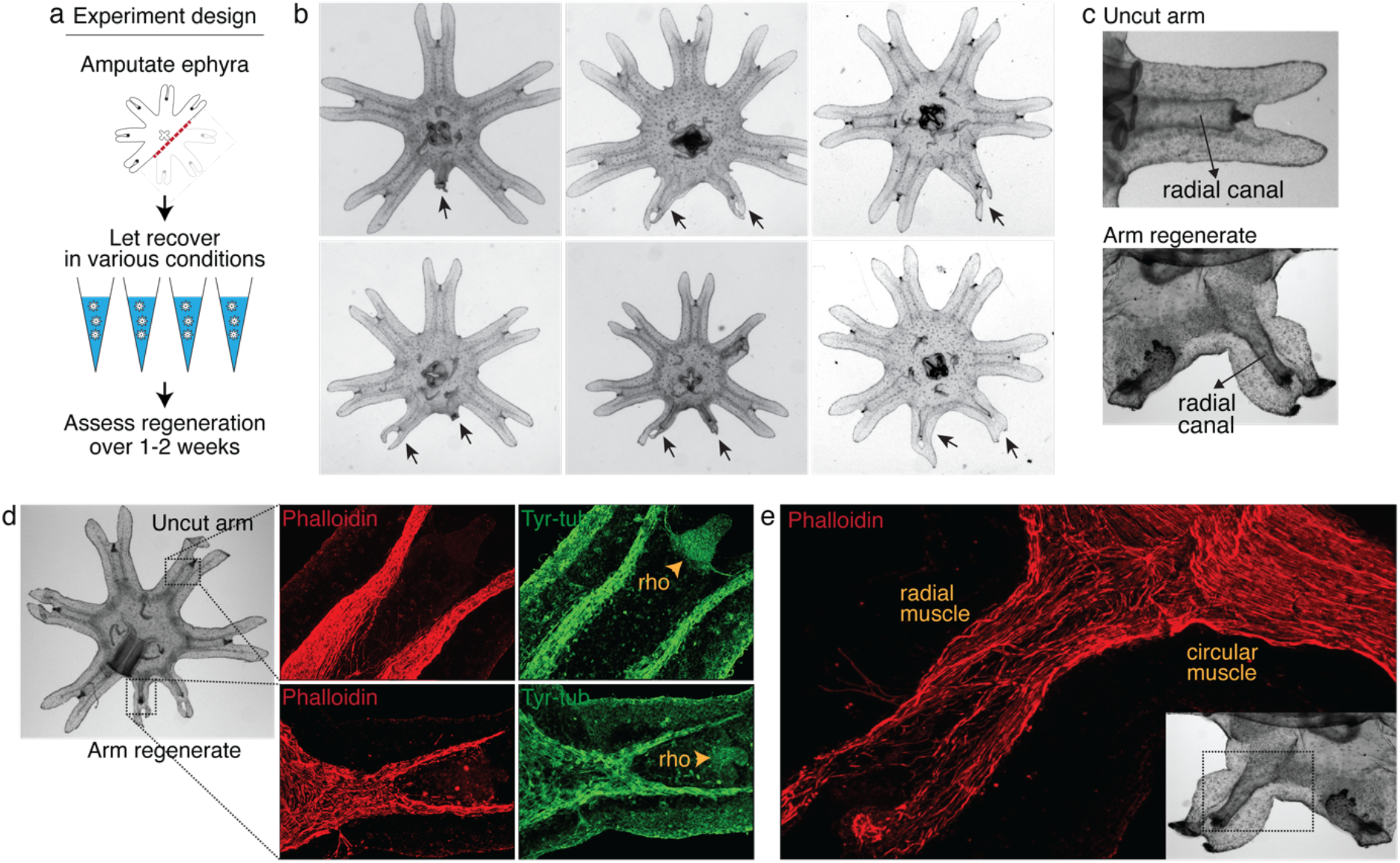
Arm regeneration in *Aurelia* ephyra can be induced using exogenous factors. **(a)** Ephyrae were amputated (red line) across the body to remove 3 arms, and then let recover in various conditions. Figure S11 tabulates the molecular and physical factors tested in the screen. Regeneration was assessed over 1-2 weeks until bell tissues began developing between the arms and obscured scoring. **(b)** Arm regeneration (arrows; from high food condition, see Figure 3a). **(c)** Radial circulatory canal in an uncut arm and is reformed in an arm regenerate. **(d)** Muscle (red), as indicated by phalloidin staining, and neuronal networks (green), as indicated by antibody against tyrosinated tubulin. The orange arrows indicate distal enrichment of tyrosinated-tubulin staining, which marks the sensory organ rhopalium (rho). Twenty ephyrae were examined and representative images are shown. **(e)** Higher magnification of the phalloidin staining shows the striated morphology of the regrown muscle in the arm regenerate (called radial muscle), which extends seamlessly from circular muscle in the body. 3 supplements: Figure S1-3

**Figure 3.**
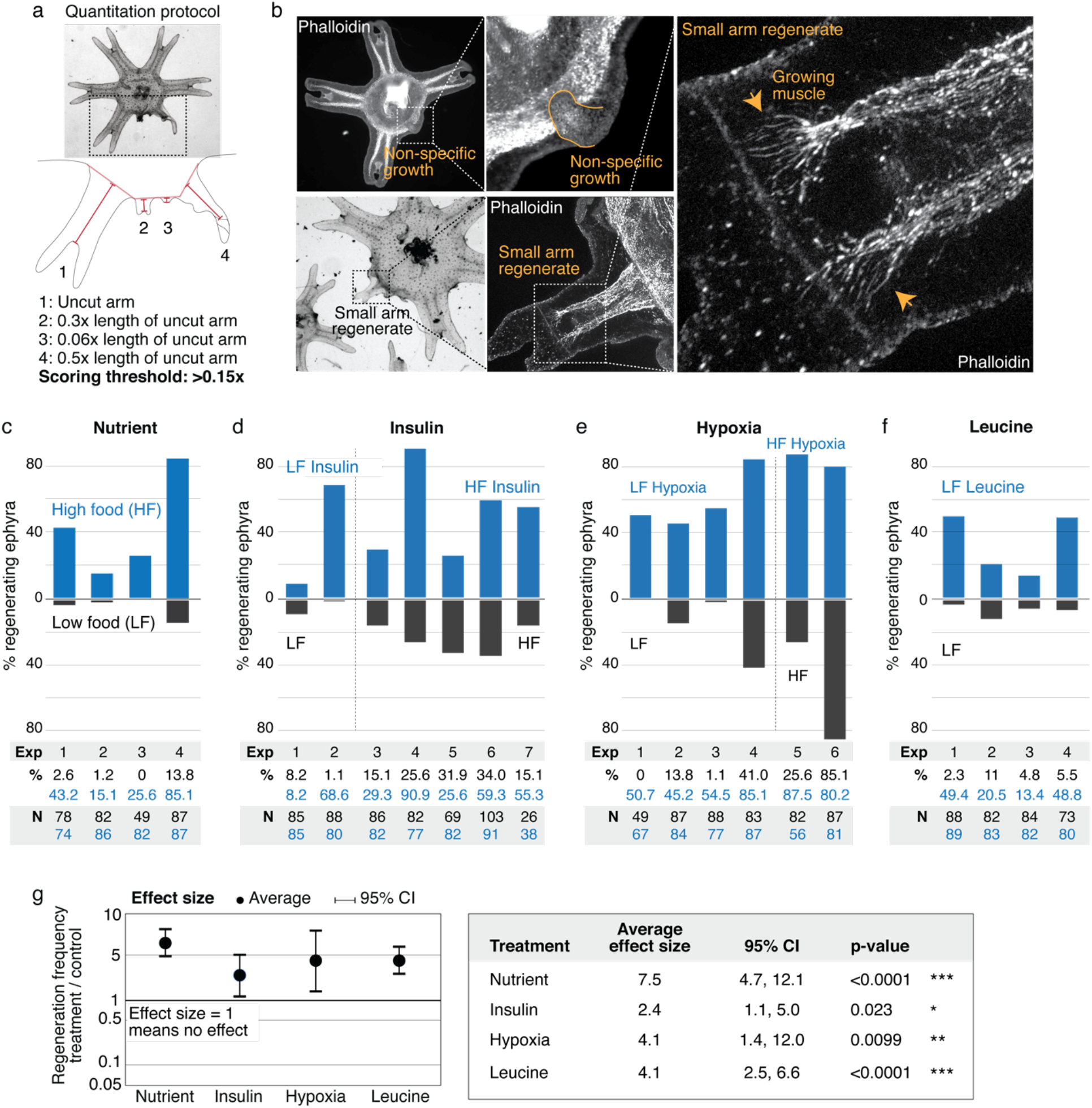
Nutrient level, insulin, hypoxia, and leucine increased regeneration frequency in *Aurelia* ephyra. **(a)** An ephyra is regenerating if it has at least one growth from the cut site with a length greater than 0.15 of the uncut arm length. The uncut arm length was determined in each ephyra by measuring 3 uncut arms and taking the average. Lappets, the distal paired flaps, were excluded in the length measurement because their shapes tend to vary across ephyrae. The measurements were performed in ImageJ. **(b)** The threshold 0.15 was chosen to balance excluding non-specific growths that show no morphological structures (*e.g.*, as shown, lack of phalloidin-stained structures) and retaining rudimentary arms that show morphological structures, including radial muscle sometime with growing ends (shown, phalloidin stained). **(c-f)** In each experiment, treated (blue) and control (grey) ephyrae came from the same strobilation. **(c)** Regeneration frequency in lower amount of food (LF) and higher amount of food (HF). The designation “high” and “low” is for simplicity, and does not presume the nutrient level in the wild. If we were to speculate, the LF amount is likely closer to typical nutrient level in the wild, based on two lines of evidence. First, regeneration frequency in LF is comparable to that observed in the natural habitat experiment. Second, in many of the wild populations studied, ephyrae mature to medusae over 1-3 months (Lucas, 2001), comparable to the growth rate in LF (by contrast, ephyrae in HF mature to medusae over 3-4 weeks). **(d)** Regeneration frequency in 500 nM insulin. **(e)** Regeneration frequency in ASW with reduced oxygen. **(f)** Ephyrae recovering in low food, with or without 100 mM L-leucine. **(g)** The effect size of a treatment was computed from the ratio between regeneration frequency in treated and control group within an experiment, *i.e.*, the metric Risk Ratio (RR; RR =1 means the treatment has no effect [Borenstein et al., 2009]). The statistical significance and reproducibility of a treatment was assessed by analyzing the effect size across experiments using the meta-analysis package, metafor (Viechtbauer 2010), in R with statistical coefficients based on normal distribution. See **Methods** for more details. A treatment was deemed reproducible if the 95% confidence intervals (95% CI) of RR exclude 1. The p-value evaluates the null hypothesis that the estimate RR is 1. Reproducibility and statistical significance of each treatment were verified using another common size effect metric, Odds Ratio (Figure S6). 6 supplements: Figure S4-S9

Intriguingly, in a few symmetrizing ephyrae, a small bud would appear at the amputation site. To follow this hunch, we repeated the experiment in the original habitat of our lab’s polyp population, off the coast of Long Beach, CA (Methods). Two weeks after amputation, most ephyrae indeed symmetrized, but in 2 of 18 animals a small arm grew (Figure 1e). This observation suggests that, despite symmetrization being the more robust response to injury, an inherent ability to regenerate arm is present and can be naturally manifest. The inherent arm regeneration presents an opportunity: Can arm regeneration be reproduced in the lab, as a way to identify factors that promote regenerative state?

To answer this question, we screened various molecular and physical factors (Figure 2a, Figure S1). Molecularly, we tested modulators of developmental signaling pathways as well as physiological pathways such as metabolism, stress response, immune and inflammatory response. Physically, we explored environmental parameters, such as temperature, oxygen level, and water current. Amputation was performed across the central body removing 3 arms (Figure 2a). Parameter changes were implemented or molecular modulators (*e.g.*, peptides, small molecules) were introduced into the water immediately after amputation. Regenerative response was assessed for 1-2 weeks until the onset of bell growth, which hindered the scoring of arm regeneration (Figure S2).

After 3 years of screening, only three factors emerged that strongly induced arm regeneration (Figure 2b). The ephyrae persistently symmetrized in the majority of conditions tested. In the few conditions where regeneration occurred, arm regenerates show multiple tissues regrown in the right locations: circulatory canals, muscle, neurons, and rhopalium (Figure 2c-e). The arm regenerates contracted synchronously with the original arms (Video 1), demonstrating a functional neuromuscular network. Thus, arm regeneration in *Aurelia* that was observed in the natural habitat can be recapitulated in the lab by administering specific exogenous factors.

The extent of arm regeneration varied, from small to almost fully sized arms (Figure 2b). The variation manifested even within individuals: a single ephyra could grow differently sized arms. Of the three arms removed, if regeneration occurred, generally one arm regenerated (67%), occasionally 2 arms (32%), and rarely 3 arms (1%, of the 4270 total ephyrae quantified in this study). Finally, the frequency of regeneration varied across clutches, *i.e.*, strobilation cohorts. Some variability may be due to technical factors, *e.g.*, varying feed culture conditions; however, variability persisted even with the same feed batch. We verified that the variability was not entirely due to genetic differences, as it manifested across clonal populations (Figure S3). Thus, there appears to be stochasticity in the occurrence of arm regeneration in *Aurelia* and the extent to which regeneration proceeds.

What are the factors that promote arm regeneration? Notably, modulation of developmental pathways often implicated in regeneration literature (*e.g.*, Wnt, Bmp, Tgfß) did not produce effect in the screen (Figure 1). We first identified a necessary condition: water current. Behaviorally, this condition promotes swimming, while in stagnant water ephyrae tend to rest at the bottom and pulse stationarily (Figure S4 and Video 2 show the aquarium setup used to implement current). In this permissive condition, the first factor that induced regeneration is the nutrient level: increasing food amount increases the frequency of arm regeneration. To measure the regeneration frequency, we scored any regenerates with lengths greater than 15% of that of an uncut arm (Figure 3a). This threshold was chosen to predominantly exclude non-specific growths or buds that show no morphological structures (Figure 3b) while including small arm regenerates that show clear morphological features, *i.e.*, lappets, radial canal, and radial muscle sometimes showing growing ends (Figure 3b). Given the clutch-to-clutch variability, control and treatment were always performed side by side using ephyrae from the same clutch. The effect size of a treatment was assessed by computing the change in regeneration frequency relative to the internal control. Statistical significance of a treatment was assessed by evaluating the reproducibility of its effect size across independent experiments (Methods). With this measurement and statistical methodologies, we found that although the baseline regeneration frequency varied across clutches, higher food amounts reproducibly increased regeneration frequency (Figure 3c). The magnitude of the increase varied (Figure 3g, 95% CI [4.7, 12.1-fold]), but the increase was reproducible (95% CI excludes 1) and statistically significant (p-value<10^−4^).

The second factor that promotes regeneration is insulin (Figure 3d). We verified that the insulin receptor is conserved in *Aurelia* (Figure S5). Administering insulin led to a reproducible (Figure 3g, 95% CI [1.1, 5.0-fold]) and statistically significant (p-value<0.05) increase in regeneration frequency. The insulin effect was unlikely to be due to non-specific addition of proteins, since bovine serum albumin at the same molarity showed no effect. Finally, the third promoter of regeneration is hypoxia (Figure 3e). We verified that the ancient oxygen sensor HIFα is present in *Aurelia* (Figure S5). Hypoxia led to a reproducible (Figure 3g, 95% CI [1.4, 12.0-fold]) and statistically significant (p-value<0.01) increase in regeneration frequency. To reduce oxygen, nitrogen was flown into the seawater, achieving ~50% reduction in dissolved oxygen level (Methods). We verified that the effect was due to reduced oxygen rather than increased nitrogen, since reducing oxygen using argon flow similarly increased regeneration frequency (95% CI [1.99, 3.3-fold], N=2 experiments, 335 ephyrae, p-value<10^−4^). The factors can act synergistically (*e.g.*, insulin and high nutrient level), but the effect appears to eventually saturate (*e.g.*, hypoxia and high nutrient level).

In addition to quantifying the number of ephyrae that regenerate, we further quantified the regeneration phenotypes in each ephyra, *i.e.*, the number of arms regenerating, the length of arm regenerates, and the formation of rhopalia (Figure S7 and S8). Nutrient level strikingly improved all phenotypic metrics: not only more ephyrae regenerated in higher nutrients, more ephyrae regenerated multiple arms, longer arms, and arms with rhopalia. Insulin and hypoxia, interestingly, show differential phenotypes. Most strikingly, while insulin induced more ephyrae to regenerate multiple arms, hypoxia induced largely single-arm regenerates, *e.g.*, hypoxia experiments 3 and 5 in Figure S7. Thus, while all factors increased the probability to regenerate, they had differential effects on the regeneration phenotypes, suggesting a decoupling to a certain extent between the regulation of the decision to regenerate and the regulation of the subsequent morphogenesis.

Of the three factors identified in the screen, nutrient input is the broadest, and prompted us to search if a more specific nutritional component could capture the effects of full nutrients in promoting regeneration. Jellyfish are carnivorous and eat protein-rich diets of zooplanktons and other smaller jellyfish (Graham and Kroutil, 2001). Notably, all three factors induced growth: treated ephyrae are larger than control ephyrae (Figure S9). The growth effect is interesting because of essential amino acids that must be obtained from food, branched amino acids supplementation correlates positively with protein synthesis and growth, and in particular, L-leucine appears to recapitulate most of the anabolic effects of high amino acid diet (Lynch and Adams, 2001; Stipanuk, 2007). Motivated by the correlation between growth and increased regeneration frequency, we wondered if leucine administration could induce regeneration. Animals typically have a poor ability to metabolize leucine, such that the extracellular concentrations of leucine fluctuate with dietary consumption (Wolfson et al., 2016). As a consequence, dietary leucine directly influences cellular metabolism. Feeding amputated ephyrae with leucine indeed led to increased growth (Figure S9). Assessing arm regeneration in the leucine-supplemented ephyrae, we observed a significant increase in the regeneration frequency (Figure 3f-g, 95% CI [2.5, 6.6-fold], p-value<10^−4^). Furthermore, leucine treatment phenocopies the effect of high nutrients, improving all measured phenotypic metrics: increasing multi-arm regeneration, the length of arm regenerate, and the frequency of rhopalia formation (Figure S7 and S8).

These experiments demonstrate that abundant nutrients, the growth factor insulin, reduced oxygen level, and the amino acid L-leucine promote appendage regeneration in *Aurelia* ephyra. The identified factors are fundamental physiological factors across animals. Might the same factors promote appendage regeneration in other animal species?

### Leucine and insulin induce regeneration in *Drosophila* limb

To pursue this question, we searched for other poorly regenerating systems, which fortunately include most laboratory models. *Drosophila*, along with beetles and butterflies, belong to the holometabolans—a vast group of insects that undergo complete metamorphosis, and that as whole, do not regenerate limbs or other appendages as adults (Hopkins and Das, 2015). Larval stages have imaginal disks, undifferentiated precursors of adult appendages such as the legs and antennae, and portions of imaginal disks have been shown to regenerate (Worley et al., 2012). Motivated by findings in *Aurelia*, we asked if leucine and insulin administration can induce regenerative response in the limb of adult *Drosophila*. We focused on testing leucine and insulin in this study because of considerations of specificity (*i.e.*, nutrients are broad and composition of nutritional needs vary across species), pragmatism (*i.e.*, administering hypoxia requires more complex setups), and in the case of *Drosophila* specifically, *Drosophila* being resistant to hypoxia (Haddad et al., 1997).

We amputated *Drosophila* on the hindlimb, across the fourth segment of the leg, the tibia (Figure 4a). The amputation removed the distal half to third of the tibia and all tarsal segments (Figure 4b). After amputation, flies were housed in vials with standard food (control) or standard food supplemented with leucine and insulin, with glutamine to promote leucine uptake (Nicklin et al., 2009) (treated) (Figure 4c). Each vial was examined multiple times, at 1, 3, 7, 14, and 21 days post amputation (dpa). Any contamination (*e.g.*, flies with uncut tibias or wrong cuts), if any, was removed at 1 and 3 dpa. Regeneration was assessed between 7-21 dpa as the presence of a regrown tibia with a reformed distal joint (Figure 4d).

**Figure 4.**
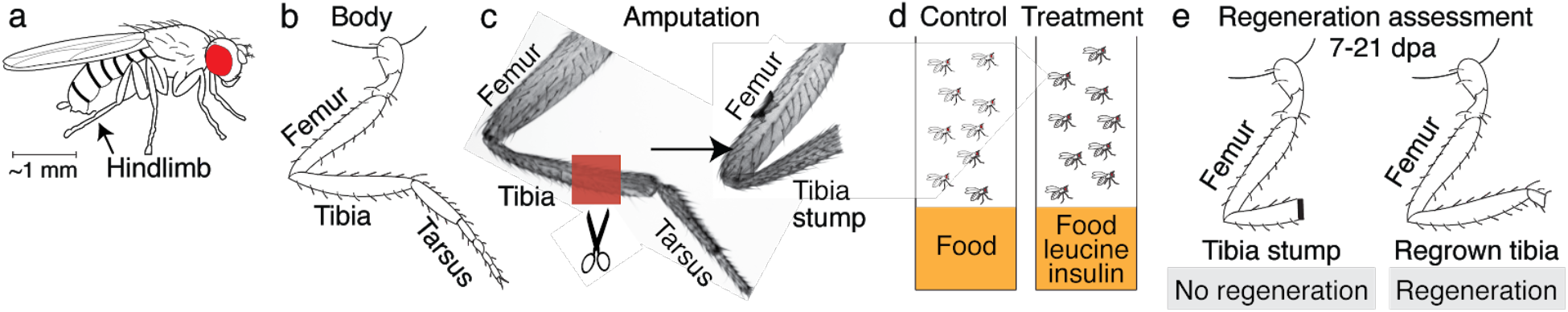
Experimental design to assess regeneration in *Drosophila* limb. **(a)** Adult *Drosophila*. **(b)** The *Drosophila* limb is a jointed limb, with rigid segments connected by flexible joints. Amputation was performed on the fourth segment, the tibia. **(c)** A hindlimb before (left) and immediately after (right) amputation. The red-shaded region indicates the amputation site. **(d)** After amputation, flies were housed in vials containing standard lab food (control) or standard lab food supplemented L-leucine and insulin (treated). **(e)** Regeneration was assessed at 7-21 days post amputation (dpa).

No regrown tibia was found in the 925 control flies examined (Figure 5a). Tibia stumps in the control flies showed melanized clots within 1-3 dpa (Figure 5b), as expected from normal wound healing process (Ramet et al., 2002), and remained so at 7-21 dpa. In the treated flies, by contrast, some amputated tibias showed no clot at 3 dpa (Figure 5c). The unclotted tips show white-colored tissues that stain positively with DAPI, indicating cellular materials, while clotted tips showed no DAPI signal (Figure 5f-h). Flies with unclotted tibia stumps were moved into a separate housing. In this population, at 7-21 dpa, a few regrown tibias were observed (Figure 5a, e). The regrown tibias culminate in reformed joints, articulating from which appears to be the beginning of a next segment. Induction of regenerative response in tibia was reproducible across genetic backgrounds, in Oregon R (12.1% white-tip tibia, 1.0% regrown tibia, N=387) and Canton S wild-type strains (29.9% white-tip tibia, 1.1% regrown tibia, N=284). Reminiscent of *Aurelia*, not all regenerative response was patterned, some flies showed non-specific outgrowth (Figure 5e).

**Figure 5.**
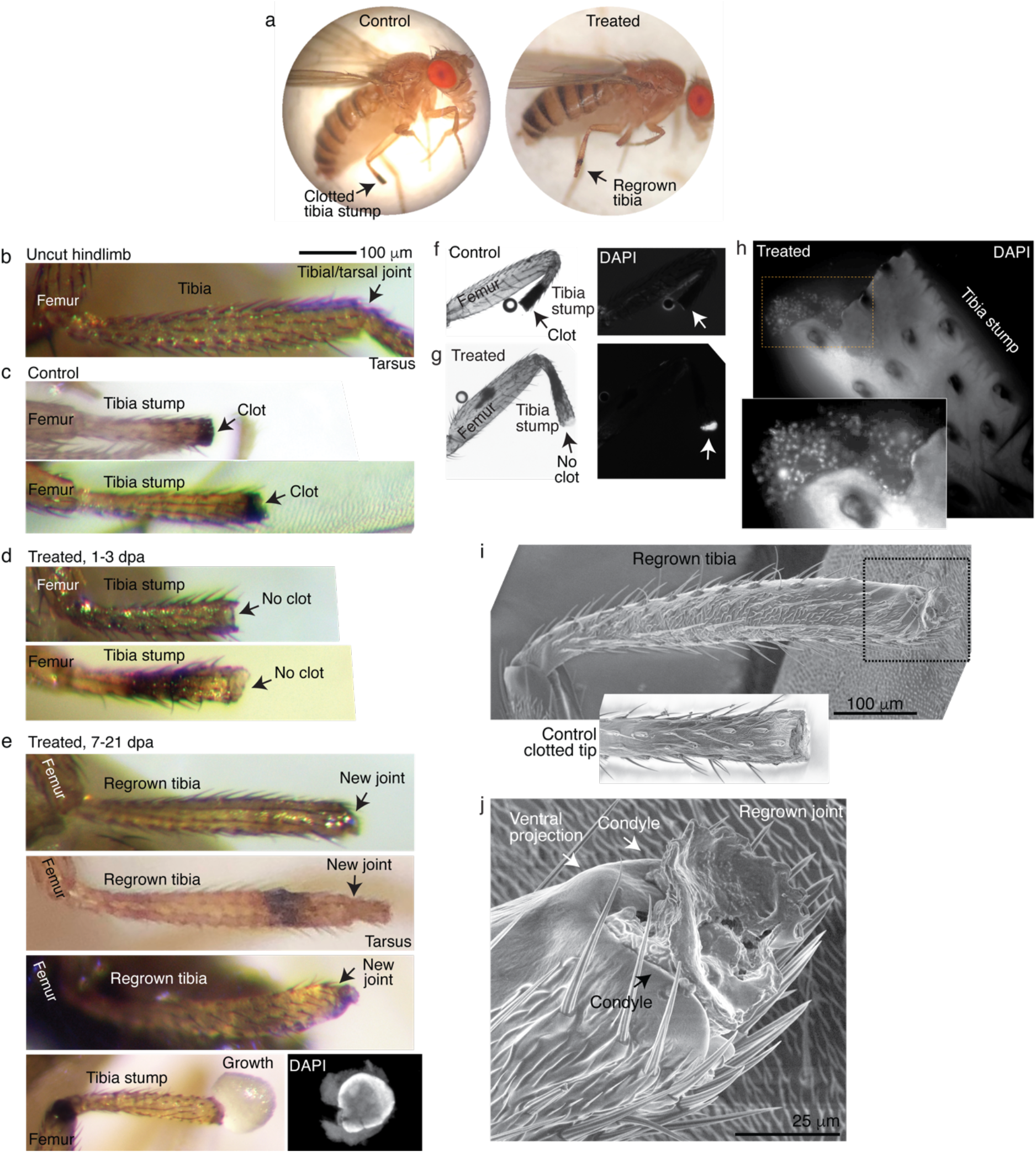
Leucine and insulin induced regeneration in *Drosophila* limb. In these experiments, upon amputation described in Figure 4, flies were placed in vials with standard laboratory food (control) or standard lab food added with 5 mM L-Leucine, 5 mM L-Glutamine, and 0.1 mg/mL insulin (treated). Doses were determined through observing the highest order of magnitude dose of amino acid that could be fed to flies over a prolonged period without shortening their lifespan. The flies were then examined at 1, 3, 7, 14, and 21 days post amputation (dpa). Color images in this figure were taken from anesthetized live flies, whereas black-and-white and fluorescent images were from dissected hindlimbs. **(a)** A control and a treated fly, imaged at 7 dpa. **(b)** An uncut hindlimb, showing distal part of femur, tibia, and proximal part of tarsus. **(c)** Control tibia stumps show melanized clotted ends from 3 dpa onward. **(d)** At 1-3 dpa, some tibia stumps in the treated population showed no clots. Sometimes a dark bruising appears near the amputation plane. **(e)** At 7-21 dpa, regrown tibias, which culminate in joints, were observed in the treated population. A dark bruise is present in one of the regrown tibias, suggesting where the amputation was. Also observed at 7-21 dpa in the treated population are some tibias stumps with non-specific growth, which stain positive for DAPI (staining method described next). **(f-g)** Tibia stumps at 3-14 dpa were dissected, fixed, and mounted in Vectashield mounting medium with DAPI. Samples from 14 dpa are shown here. Insect cuticle is not dissected to restrict DAPI penetrance only to the distal tip. Clotted tips of control tibia stumps did not stain with DAPI (**f**, 10 of 10), whereas unclotted tips of treated tibia stumps stained with DAPI (**g**, 14 of 16). **(h)** Higher-resolution confocal image of an unclotted tip of a treated tibia stump at 14 dpa showing DAPI-positive cells. **(i)** Fly with a regrown tibia at 21 dpa (an earlier picture of this regrown tibia is the top panel in Figure e) was mounted onto an environmental SEM with a copper stub. Inset shows a clotted tibia stump from a control fly, with the discoloration at the end corresponding to the clot. **(j)** Magnification of the regenerated joint, with the arrows denoting the two condyles and the additional ventral projection.

Scanning electron micrograph of a regrown tibia (the top tibia in Figure 5e, taken one week later) morphologically confirms the regenerated joint as a tibial/tarsal joint. The completed tibia is enclosed in a sclerotized cuticle lined with longitudinal arrays of bristles, with no visible signs of the previous amputation (Figure 5i). The joint-like structure shows the expected bilateral symmetry of a tibial/tarsal joint (as opposed to *e.g.*, the radially symmetrical tarsal/tarsal joint) (Mirth and Akam, 2002)with rounded projections at the posterior and anterior end (arrows in Figure 5j). These projections, called condyles, function as points of articulation between opposing leg segments. Indeed, articulating from the regrown condyles appears to be further growth. Finally, a unique feature of the tibial/tarsal joint of the hindlimb (but not of fore or midlimb) is an additional ventral projection between the side condyles (Mirth and Akam, 2002), which serves to restrain bending of the leg upward. The ventral projection is indeed present in the regenerated joint (arrow in Figure 5j).

### Leucine and sucrose induce regeneration in mouse digit

The ability of leucine and insulin to induce regenerative response in *Drosophila* limb and *Aurelia* appendage motivated testing in vertebrates. One sign that limb regeneration may be feasible in humans is that fingertips regenerate (Illingworth, 1974). The mammalian model for studying limb regeneration is the house mouse, *Mus musculus*, which like humans regenerates digit tips. Although more proximal regions of digits do not regenerate, increasing evidence suggests that they have inherent regenerative capacity. In adult mice, implanting developmental signals in amputated digits led to specific tissue induction, *i.e.*, bone growth with Bmp4 or joint-like structure with Bmp9 (Yu et al., 2019). In neonates, reactivation of the embryonic gene *lin28* led to distal phalange regrowth (Ng et al., 2013). Thus, while patterned phalange regeneration can be induced in newborns, induction in adults so far involves a more fine-tuned stimulation, *e.g.*, to elongate bone and then make joint, Bmp4 was first administered followed by Bmp9 in a timed manner. Motivated by the findings in *Aurelia* and *Drosophila*, we tested if leucine and insulin administration could induce a more self-organized regeneration in adult mice.

We performed amputation on the hindpaw (Figure 6a), on digit 2 and 4, leaving the middle digit 3 as an internal control (Figure 6b). To perform non-regenerating amputation, a clear morphological marker is the nail, which is associated with the distal phalange (P3). Amputation that removes <30% of P3 length, that cuts within the nail, readily regenerates, whereas amputation that removes >60% of P3 length, corresponding to removing almost the entire visible nail, does not regenerate (Figure 6c) (Chamberlain et al., 2017; Lehoczky et al., 2011). We therefore performed amputations entirely proximal to the visible nail – giving, within the precision of our amputation, a range of cut across somewhere between the proximal P3 and the distal middle phalange (P2) (Figure 6d) – a range that is well below the regenerating tip region. Note additional morphological markers that lie within the non-regenerating region: the os hole (‘o’ in Figure 6c), where vasculatures and nerves enter P3, the bone marrow cavity (‘bm’ in Figure 6c), and the sesamoid bone (‘s’ in Figure 6c) adjacent to P2.

**Figure 6.**
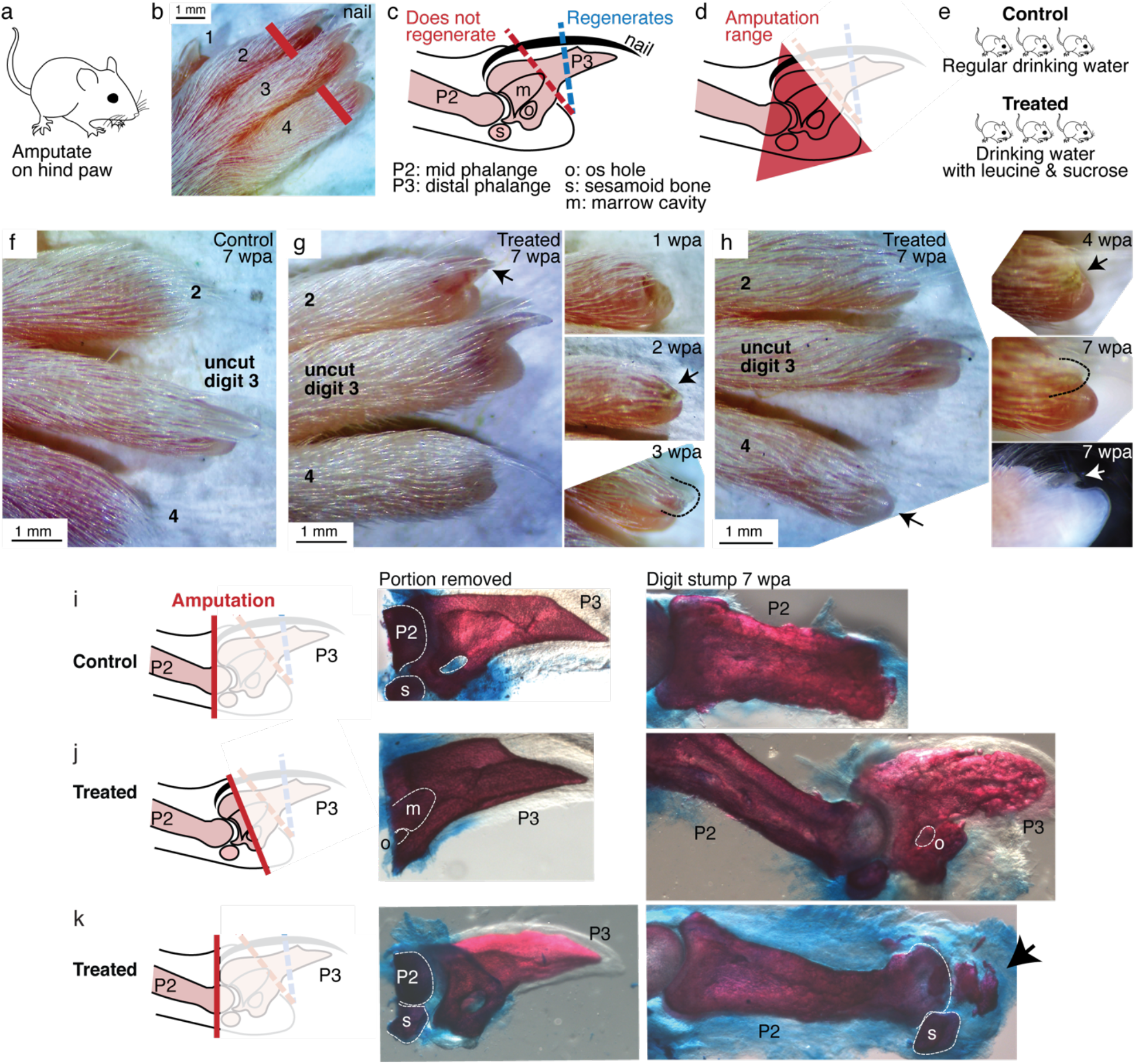
Leucine and sucrose induced regeneration in adult mouse digit. **(a-b)** Amputation was performed on hindpaws of adult (3-6 month old) mice, on digits 2 and 4, proximal to the nail. **(c)** Schematic of the distal phalange (P3) and middle phalange (P2). Amputations that remove <30% of P3 (blue line) regenerate, whereas amputations that remove >60% of P3 (red line) do not regenerate. Amputations in the intermediate region can occasionally show partial regenerative response. **(d)** Amputations in this study were performed within the red-shaded triangle. **(e)** Amputated mice were given regular drinking water (control) or drinking water supplemented with 1.5% L-leucine, 1.5% L-glutamine, and 4-10 w/v % sucrose (2 exps with 4%, 6 exps with 10%). Drinking water, control and treated, was refreshed weekly. **(f)** A representative paw from the control group. The amputated digits 2 and 4 simply healed the wound and did not regrow the distal phalange. **(g)** In this treated mouse, digit 2 (arrow) regrew the distal phalange and nail. Insets on the right show the digit at earlier time points. At week 1, the amputation site still appeared inflamed. At week 3, the beginning of the nail appears (arrow). At week 3, a clear nail plate was observed. **(h)** In this treated mouse, digit 4 (arrow) regrew and began to show nail reformation by week 4 (top inset, see arrow), that turns into a clear nail plate by week 7 (middle inset), as can be seen more clearly from the side-view darkfield image (bottom inset). **(i-k)** Whole-mount skeletal staining. Dissected digits were stained with Alizarin red, an anionic dye that highly localizes to the bone. Left panels show illustration of the amputation plane, middle panels show skeletal staining of the portions removed, and right panels show skeletal staining of the digit stumps 7 weeks after amputation. 2 supplements: Figure S10 and S11

The digit portion removed was immediately fixed for control. The amputated mice were either provided with water as usual (control) or water supplemented with leucine and sucrose (treated) (Figure 6e). Both groups were monitored for 7 weeks. Sucrose was used because insulin is proteolytically digested in the mammalian gut. The sucrose doses used are lower or the administration duration is shorter than those shown to induce insulin resistance (Cao et al., 2007; Togo et al., 2019). We verified that control and treated mice had comparable initial weights (35.1±0.6 vs 34.1±1.1 grams, p-value=0.402, student’s t-test), and that as expected from amino acid and sugar supplementation, treated mice gained more weight over the experimental duration (4.5±1.0 vs 7.8±1.0 grams, p-value=0.028, student’s t-test).

As expected for amputation proximal to the nail, no regeneration was observed in the control mice (N=20 digits, 10 mice). Amputated digits healed and re-epithelialized the wound as expected (Figure 6f). Skeletal staining shows blunt-ended digit stumps (Figure 6i) and in many instances, as expected, dramatic histolysis, a phenomenon where bone recedes further from the amputation plane (Figure S10) (Chamberlain et al., 2017). By contrast, 18.8% of the treated digits (N=48 digits, 24 mice) showed various extents of regenerative response (Figure S10).

We observed, as in *Aurelia* and *Drosophila*, an unpatterned response (Figure S10), wherein skeletal staining reveals excessive bone mass around the digit stump, similarly to what was observed in some cases with BMP stimulation (Yu et al., 2019). However, we also observed patterned responses (Figure S11). The most dramatic regenerative response was observed in 2 digits (Figure 6g-h). In one digit, an almost complete regrowth of the distal phalange and the nail was observed (Figure 6g). Skeletal staining of the portion removed from this digit (Figure 6j) shows that it was amputated at the proximal P3 transecting the os hole. By 7 weeks, skeletal staining of the regrown digit (Figure 6j) shows that the P3 bone was almost completely regrown. The regrown P3 shows trabecular appearance that is similar in general structure but not identical to the original P3. Another dramatic response was observed from another digit, which began reforming the nail by 7 weeks (Figure 6h). Skeletal staining of the portion removed from this digit shows that it was amputated across the P2 bone, removing the entire epiphyseal cap along with the sesamoid bone (Figure 6k). Skeletal staining of the regenerating digit shows that the epiphyseal cap was regrown, along with its associated sesamoid bone. Moreover, articulating from the regenerated P2 appears to be the beginning of the next phalangeal bone (arrow, Figure 6k). To our knowledge, the regenerative response observed in these digits represents the most dramatic extent of self-organized mammalian digit regeneration reported thus far. Distal phalange regeneration in adults has not been reported, while interphalangeal joint formation from a P2 amputation has been achieved only through sequential Bmp administration (Yu et al., 2019) and there has been no documentation of the regrowth of the sesamoid bone.

## Conclusion

In this study, amputations were performed on *Aurelia* appendage, *Drosophila* limb, and mouse digit. None of these animals are known to regenerate robustly (*Aurelia*) if at all (*Drosophila* and mouse) from these amputations. Upon administration of L-leucine and sugar/insulin, dramatic regenerative response was observed in all systems. The conserved effect of nutrient supplementation across three species that span 500 million years of evolutionary divergence suggests energetic parameters as ancestral regulators of regeneration activation in animals.

While we did not test the appendage regenerative effect of hypoxia beyond *Aurelia*, it is notable that in mice hypoxia coaxes cardiomyocytes to re-enter cell cycle (Kimura et al., 2015) and activating HIFα promotes healing of ear hole punch injury (Zhang et al., 2015). The diverse physiologies of animals across phylogeny may seem difficult to reconcile with a conserved regulation of regeneration, especially in the view of regeneration as recapitulation of development. Growing a jellyfish appendage is different from building a fly leg or making a mouse digit. However, there is another way of looking at regeneration as a part of tissue plasticity (Galliot and Ghila, 2010). In this view of regeneration, before tissue-specific morphogenesis commences, a more upstream regulation is hypothesized that controls the broadly shared processes of growth, proliferation, and differentiation. In support of this idea, regeneration across species and organs relies one way or another on the presence of stem cells or differentiated cells re-entering cell cycle and re-differentiating (Cox et al., 2019). We propose that in animals that poorly regenerate, high nutrient input turns on growth and anabolic states that promote tissue rebuilding upon injury.

That regenerative response can be induced blurs the boundary between regenerating versus non-regenerating animals. The factors identified in the study are not exotic: variations in amino acids, carbohydrates, and oxygen levels are conditions that the animals can plausibly encounter in nature. These observations highlight two potential insights into regeneration. First, regeneration is environmentally dependent. An animal would stop at wound healing under low-energy conditions and regenerate in energy-replete conditions. In this view, for the animals examined in this study, the typical laboratory conditions may simply not be conducive to regeneration. Alternatively, the interpretation we favor, what we observed is dormant regeneration, which can be activated with broad environmental factors. We favor this interpretation because the regenerative response was unusually variable. The variability stands in stark contrast to the robust regeneration in *e.g.*, axolotl, planaria, or hydra. Just like mutations produce phenotypes with varying penetrance and expressivity, the variable regenerative response speaks to us as a fundamental consequence of activating a biological module that has been evolutionarily inactivated. The ordinariness of the activators suggests ancestral regeneration as part of a response to broad environmental stimuli.

In particular, the conserved effects of nutrient supplementation suggest that regeneration might have originally been a part of growth response to abundant environments. No nutrient dependence has been observed in highly regenerating animal models such as planaria, hydra, and axolotl. Environment-dependent plasticity, however, is pervasive in development, physiology, behavior, and phenology (West-Eberhard, 2003; Mockzek et al., 2011). We therefore conjecture that environment-dependent plasticity may have characterized the ancestral form of regeneration. In this conjecture, present regenerating lineages might have decoupled the linkage with environmental input and genetically assimilated regenerative response — because regeneration is adaptive or coupled to a strongly selected process, *e.g.*, reproduction. In parallel, non-or poorly regenerating animals might have also weakened the linkage with environmental input, but to silence the regenerative response. This predicts an ancient form of a robustly regenerative animal (like planaria, hydra, axolotl) that tunes its regeneration frequency to nutrient abundance. Such plasticity has been reported in the basal lineage Ctenophora (Bading et al., 2017).

In conclusion, this study suggests that an inherent ability for appendage regeneration is retained in non-regenerating animals and can be unlocked with a conserved strategy. While the observed regenerative response is not perfect, this motivates further investigation into potentially more promoting factors or the possibility of combining broad promoting factors with species- or tissue-specific morphogenetic regulators. Reiterating Spallanzani’s hope, Marcus Singer supposed half a century ago that “… every organ has the power to regrow lying latent within it, needing only the appropriate ‘useful dispositions’ to bring it out (Singer, 1958).” The surprise, in hindsight, is the simplicity by which the regenerative state can be promoted with *ad libitum* amino acid and sugar supplementation. This simplicity demonstrates a much broader possibility of organismal regeneration, and can help accelerate progress in regeneration induction across animals.

## Acknowledgements

The authors thank Kiersten Darrow and Michael Schaadt at the Cabrillo Marine Aquarium, Cabrillo, CA for the gifts of *Aurelia* polyps and help with jellyfish culture, Matthew Hunt from the Kavli Nanoscience Institute at Caltech for help with SEM imaging, James McGehee, Angela Stathopolous, and Kai Zinn for sharing *Drosophila* strains, Yujing Yang and Long Cai for help with imaging, and Carlos Lois for demonstration of digit amputation. We thank Gertrud Schupbach, Natalie Andrew, Anish Sarma, and Aki Ohdera for comments on the manuscript; and Bruce Hay, Dianne Newman, Katalina Fejes-Toth, Luis del Peso, Aditya Saxena, Joe Parker, Marta Truchado Garcia, and Justin Bois for discussion. This work was supported by the National Science Foundation Graduate Research Fellowship Program (1144469; to M.J.A.), the James E. and Charlotte Fedde Cordes Postdoctoral Fellowship in Biology (to D.A.G.); the James S. McDonnell Foundation for Complex Systems Science (220020365; to L.G.), the Center for Environmental Microbial Interactions at Caltech, and Charles Trimble and Caltech’s Biology and Biological Chair’s Council Inducing Regeneration Fund (to L.G.).

## Author contributions

M.J.A., F.T., and L.G. conceptualize the paper. M.J.A. and T.B. designed and performed experiments and data analysis in *Aurelia*. F.T. and M.H. designed and performed experiments and data analysis in *Drosophila* and mouse. Y.L designed and performed experiments and data analysis in a second *Drosophila* strain. M.R., J.O.D., and D.A.G. contributed experiments in *Aurelia* and discussion into the study. P.L. contributed help with *Aurelia* experiments in the natural habitat. M.J.A., F.T., and L.G. wrote the manuscript draft. All authors reviewed and edited the manuscript.

---

**Figure S1.**
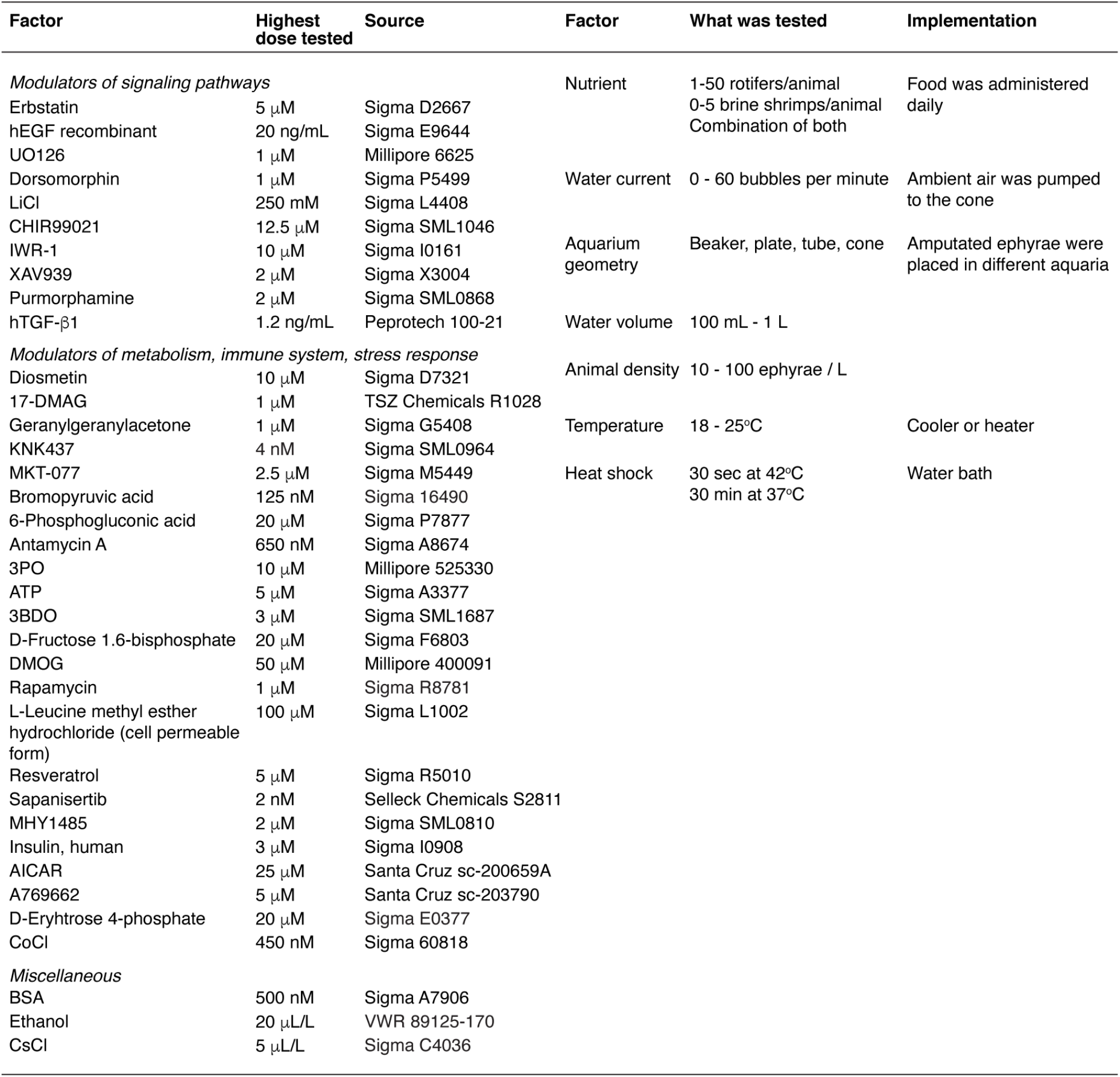
Various molecular and physical modulations were screened to recapitulate arm regeneration. Modulators were administered or physical parameters were implemented upon amputation. Some factors were dissolved in DMSO or ethanol; for these molecules, the control group was administered with an equal volume of the solvent. Since few, if at all, of the molecular modulators had been tested in *Aurelia*, the maximum concentrations were tested to maximize the chance of seeing an effect. Maximum concentration was determined by solubility in saltwater or onset of adverse effects (*e.g.,* degrowth, paralysis, death) upon overnight incubation. Where available, previously reported concentrations in cell culture or animal systems were included in the testing. A negative result means no obvious effects were observed at the maximum concentration that warrant further investigation. For factors that gave interesting effects (*e.g.*, insulin), a range of lower concentrations were subsequently tested for optimization.

**Figure S2.**
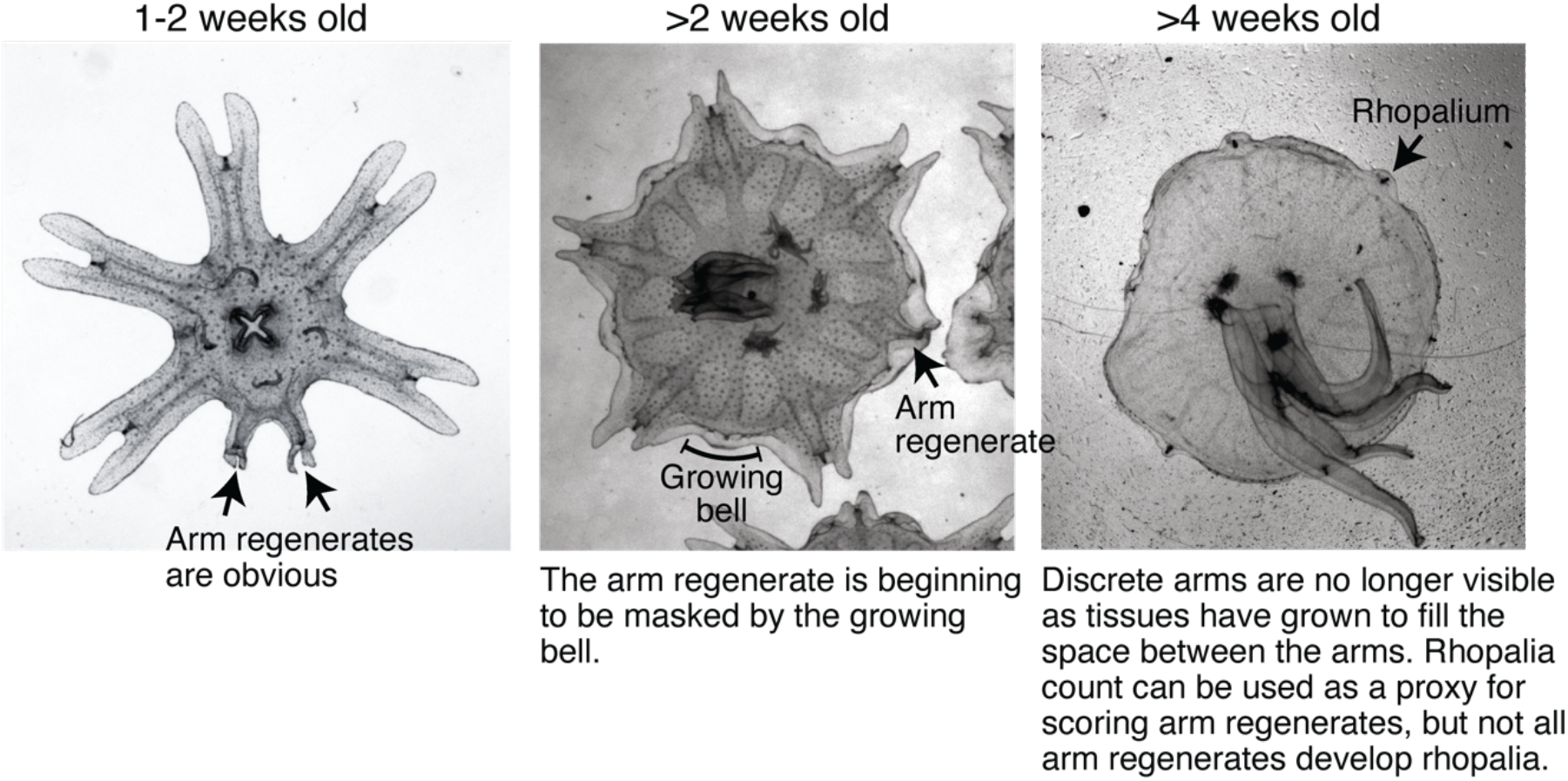
Bell growth limited the time window for assessing arm regeneration. Ephyrae in the lab mature into full-belled medusae within ~4 weeks. The transition to medusa commences at 1-2 weeks after strobilation, with the onset of bell growth. Over 2-3 weeks, body tissues gradually grow and fill between the discrete arms to form a continuous bell characteristic of a medusa. Arm regeneration can be unambiguously scored in ephyrae before the bell has significantly grown. Bell growth also limited testable doses in some factors, *e.g.*, testing higher food amounts than reported here led to accelerated bell growth at a rate that did not allow enough time window to quantify regeneration.

**Figure S3.**
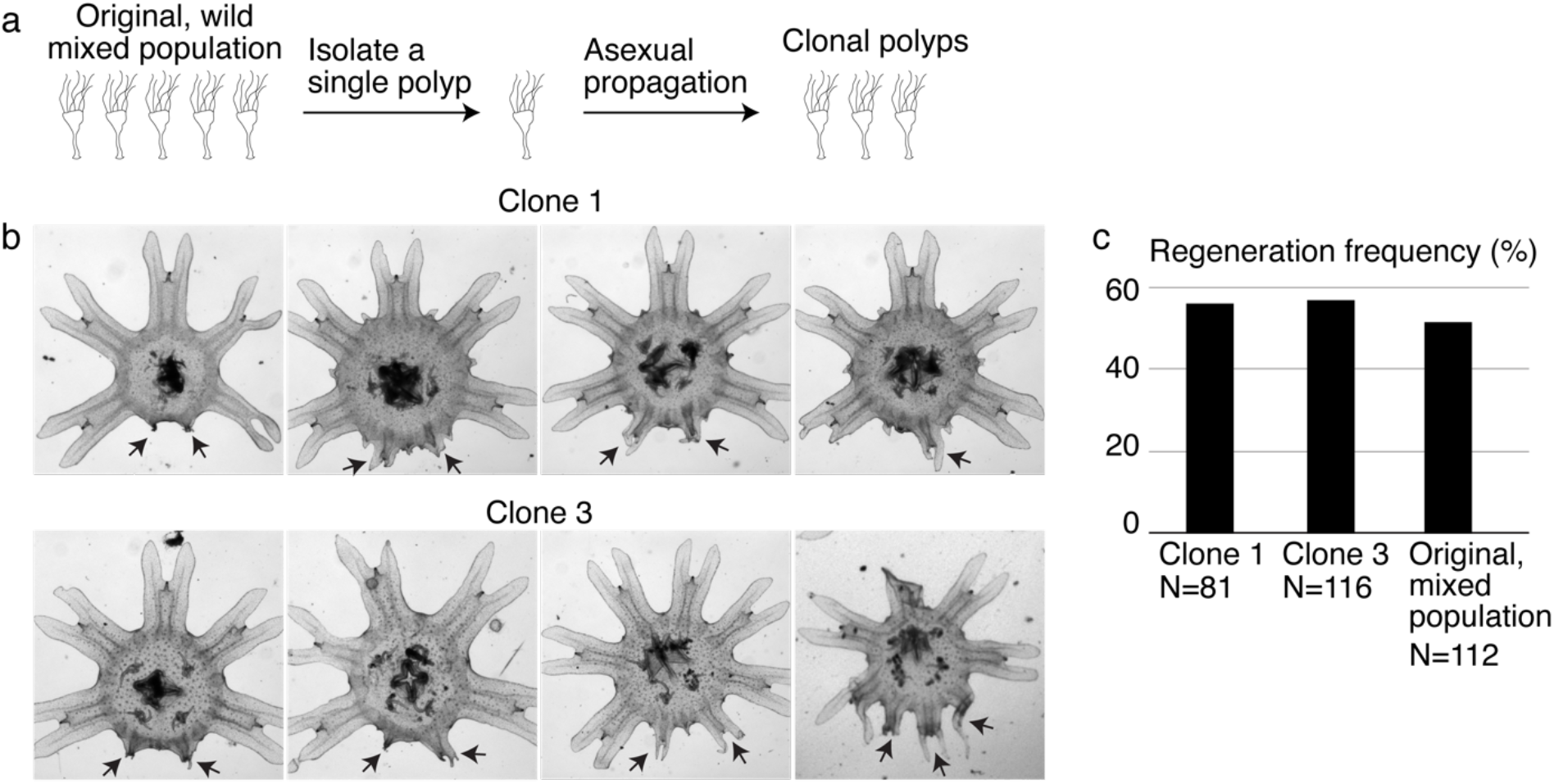
Variable extent of regeneration was observed in clonal lines. **(a)** To develop genetically clonal lines, single polyps were isolated and settled onto tissue culture dishes. Within 1-3 months, with daily feeding of enriched brine shrimps, each dish was re-populated with polyps asexually budding from the single parental polyp. **(b)** Regeneration induction with high food performed in two clonal lines. Arrows indicate arm regenerates. **(c)** Regeneration frequency in the clonal and original mixed populations measured in the same experiment. The data reported in the main text come from experiments performed in clone 3.

**Figure S4.**
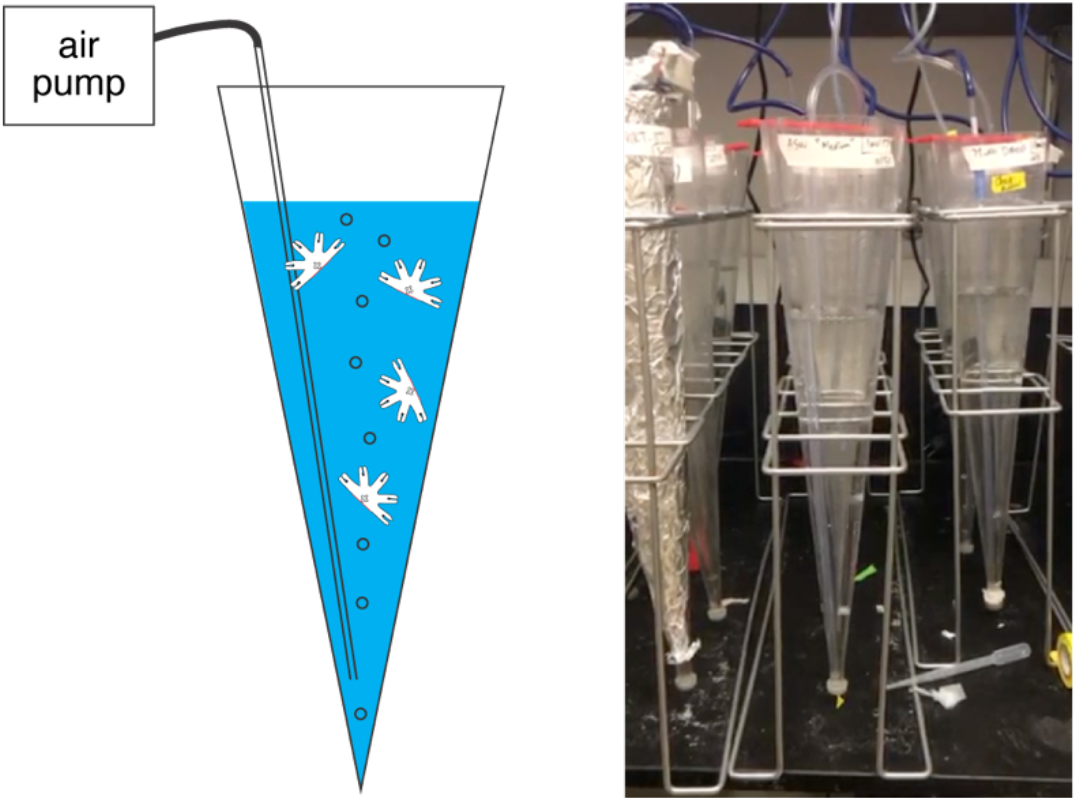
Water current is a permissive requirement for arm regeneration induction. Various physical environments for the ephyrae recovering from injury were tested, *e.g.*, shallow vs deep water, seawater with varying salinity, cold vs warm temperature, light versus dark, stagnant water vs current, generating water current through various means, including shaking or rotating to generate turbulent mixing and as shown here air bubbling a conical tube to generate vertical current (shown here). While symmetrization occurred robustly in all conditions, consistent induction of regeneration only occurred in the presence of columnar water current. The experiments presented in this study were performed in the bubbler cone setup, where a 1L sand settling cone was repurposed into an aquarium and connected to an air pump to generate a gentle current of ~1 bubble/second (**Movie S2**). In this setup, the ephyrae were continually swimming along the current, either upward along the bubble-generated current or downward along the gravity-driven current.

**Figure S5.**
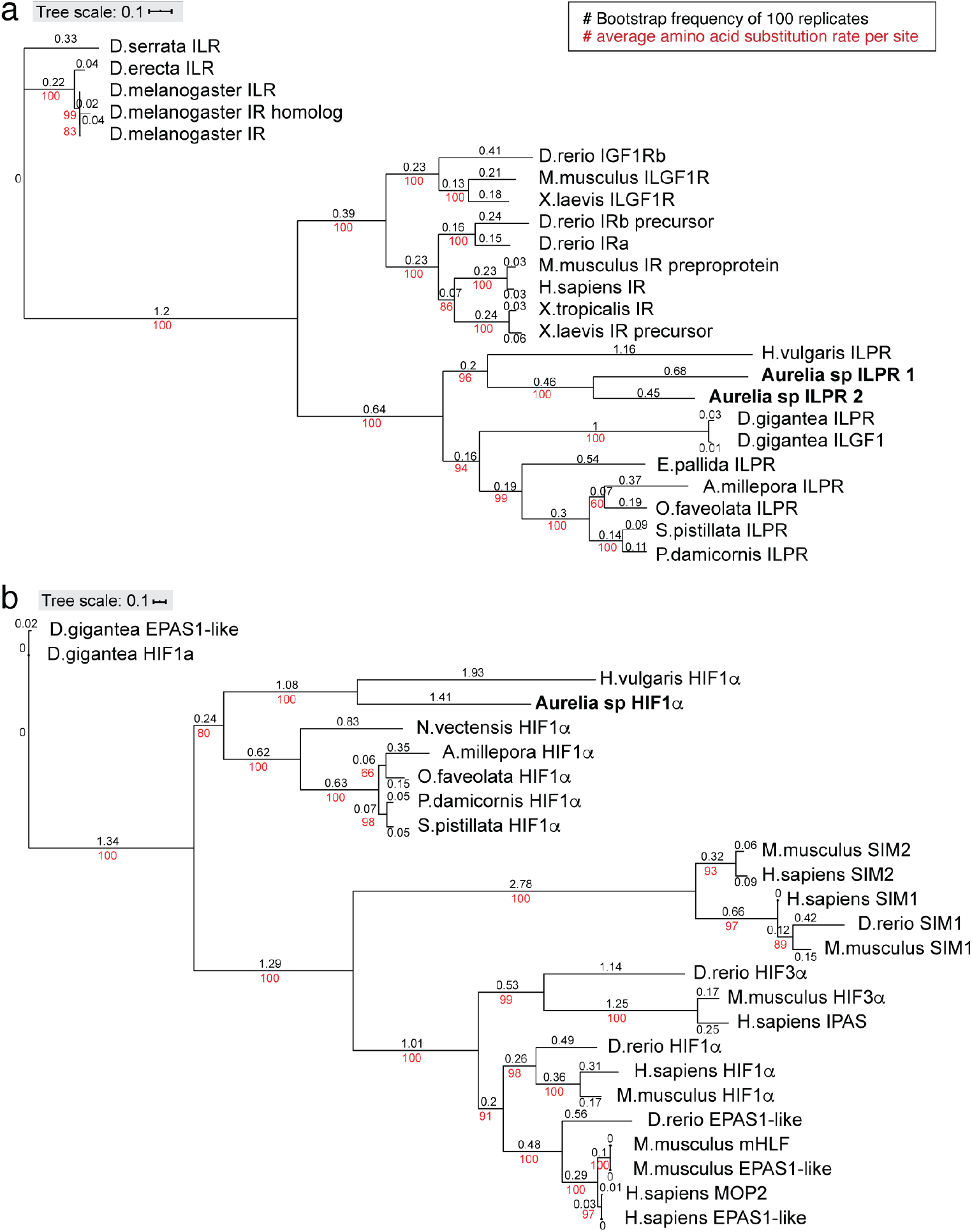
Conservation of insulin receptor and HIFα in *Aurelia*. Phylogenies of insulin receptor **(a)** and HIFα **(b)** genes were constructed using the maximum likelihood inference computed with the IQ-TREE stochastic algorithm (Nguyen et al., 2015), and visualized using ITOL (https://itol.embl.de/upload.cgi). These trees verify the he simple trees are not meant to be comprehensive, but a verification of the genes annotated as insulin-like protein receptor (ILPR) and HIFα in the *Aurelia* gene models by testing conservation with their known counterparts in other organisms. IQ-TREE parameters: Insulin receptor consensus tree is constructed from 1000 bootstrap trees; log-likelihood of consensus tree is −45374.0; the Robinson-Foulds distance between ML and consensus tree is 0. HIFα consensus tree is constructed from 1000 bootstrap trees; log-likelihood of consensus tree is −24414.4; the Robinson-Foulds distance between ML and consensus tree is 0.

**Figure S6.**
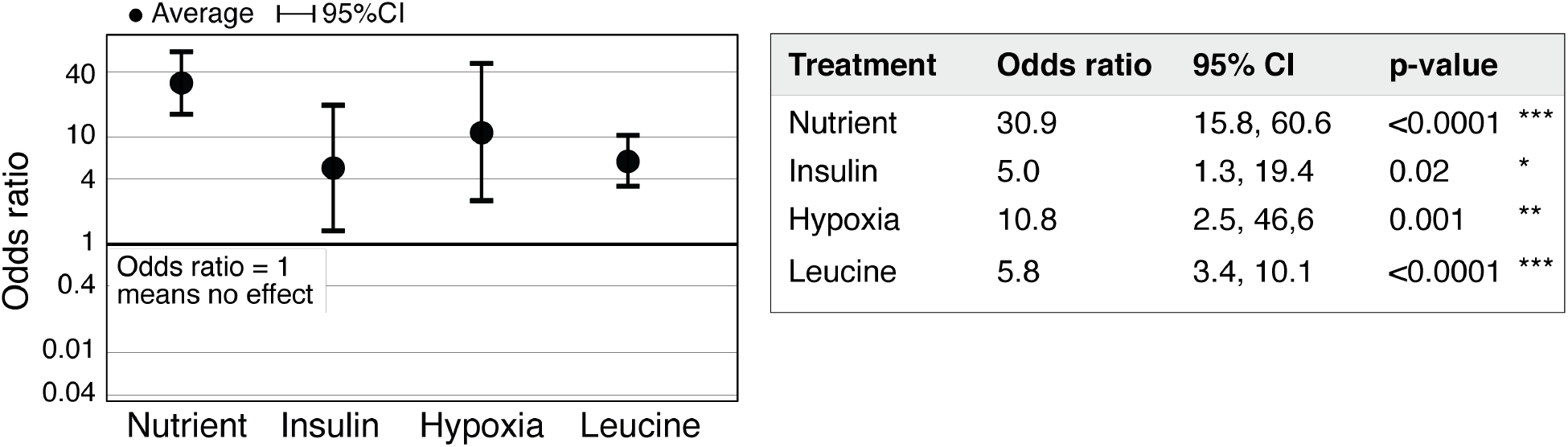
Statistical significance of regeneration induction in *Aurelia* assessed using Odds Ratio. In addition to RR analysis presented in Figure 3g, another common measure of effect size is the Odds Ratio (OR) (Borenstein et al., 2009). OR compares the odds of outcome in the presence vs. absence of treatment (Methods). Analysis of OR across experiments was performed using the metafor package (Viechtbauer, 2010) in R with statistical coefficients based on normal distribution (Methods). A treatment is reproducible if the 95% confidence intervals (95% CI) exclude 1. The p-value evaluates the null hypothesis that the estimate OR is 1.

**Figure S7.**
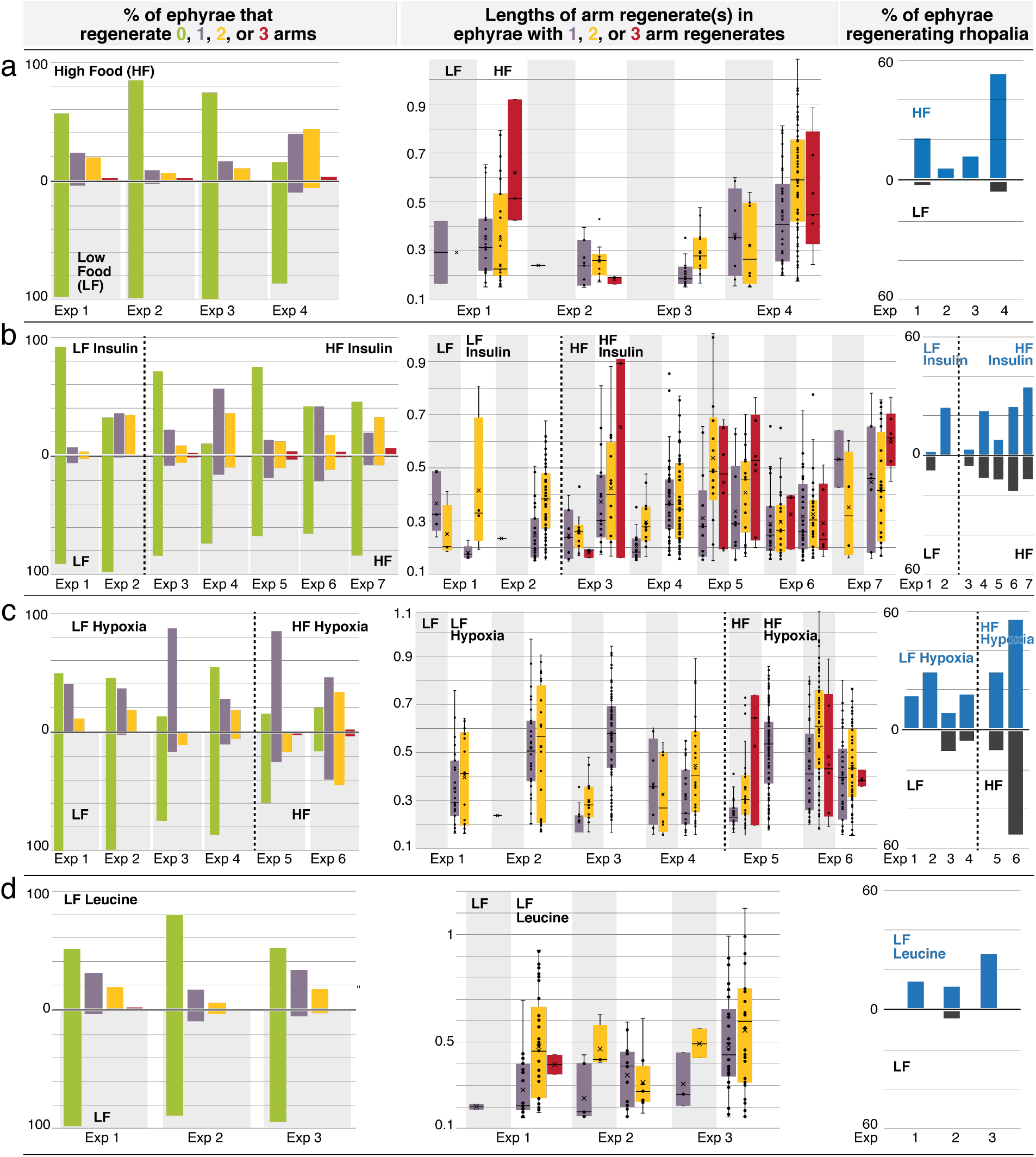
**Regeneration phenotypes** in **(a)** high amount of nutrients, **(b)** insulin, **(c)** hypoxia, and **(d)** L-leucine. For each treatment, Left: The percentage of ephyrae that regenerate 0 (green), 1 (purple), 2 (yellow), or 3 arms (red). Middle: The length(s) of arm regenerate(s) in ephyrae that regenerate 1 arm (purple), 2 arms (yellow), and 3 arms (red) – normalized to the average length of uncut arms in the same ephyra. For ephyrae with multiple arm regenerates, lengths of all arms were measured and plotted individually. Boxplot: median (line), average (cross), 1st and 3rd quartiles (the box), 5^th^ and 95^th^ percentile (whiskers), and individual data points (black circles). Right: The percentage of ephyrae that reform rhopalia in control (grey) and treated (blue) groups.

**Figure S8.**
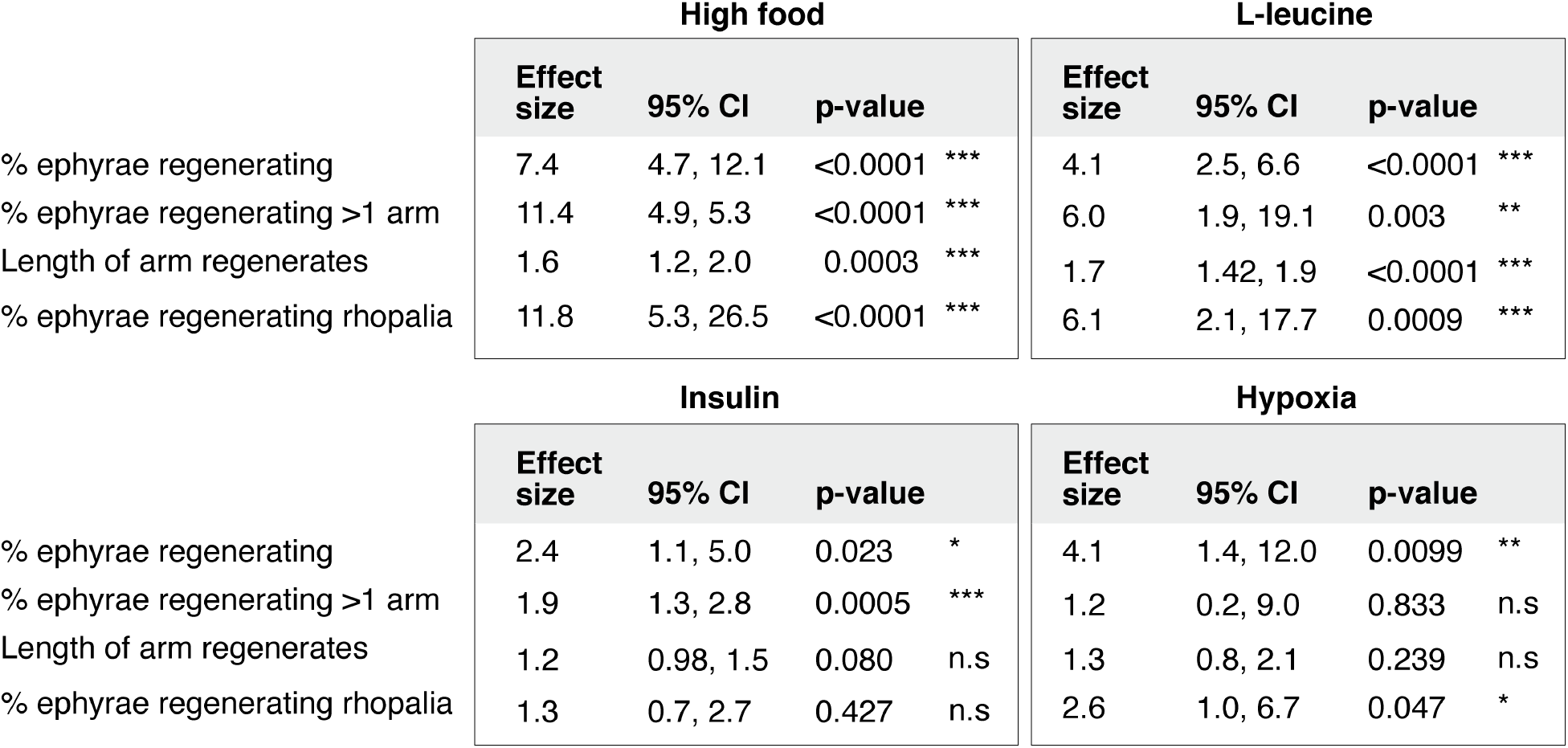
Statistical analysis of the regeneration phenotypes in high amount of nutrients, insulin, hypoxia, and L-leucine. For frequency measurements, the effect size of a treatment compares the probability of an outcome in treated vs. control group (i.e., Risk Ratio, Methods). For length measurement, the effect size of a treatment compares the proportionate change that results from the treatment (i.e., Response Ratio, Methods). Analysis of effect size across experiments was performed using the metafor package15 in R with statistical coefficients based on normal distribution (Methods). A treatment is reproducible if the 95% confidence intervals (95% CI) exclude 1. The p-value evaluates the null hypothesis that the estimate effect size is 1 (i.e., no effect).

**Figure S9.**
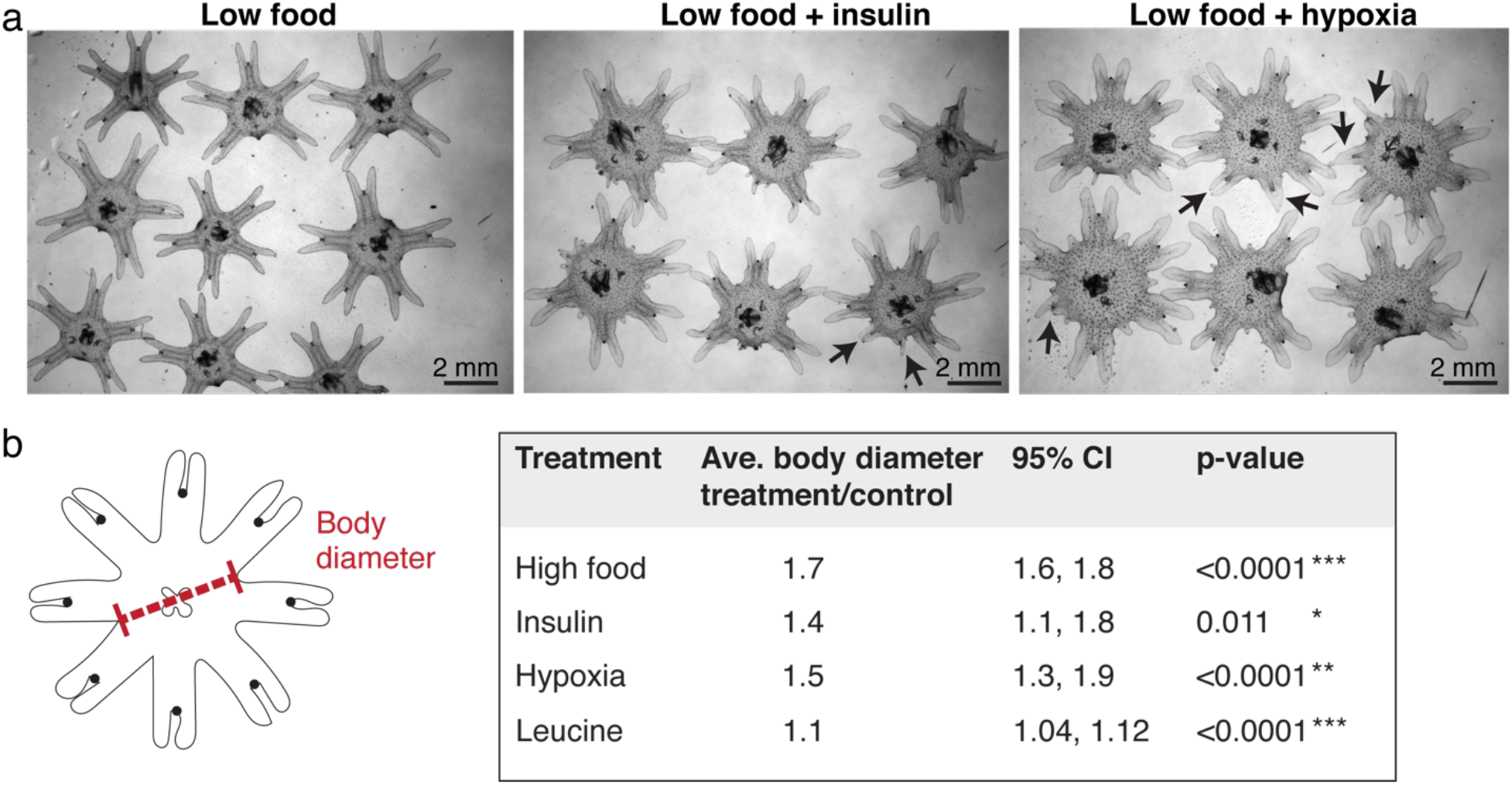
Ephyrae in high food, insulin, or hypoxia, and L-leucine tend to be bigger in size. **(a)** Representative images of ephyrae growing in low food, 500 nM insulin, and hypoxia. Black arrows indicate regenerating arms. **(b)** Effect size analysis of the body size increase was performed using the metafor package (Viechtbauer et al., 2010)in R (Methods). A treatment effect is reproducible if the 95% CI exclude 1. The p-value evaluates the hypothesis that there is no effect.

**Figure S10.**
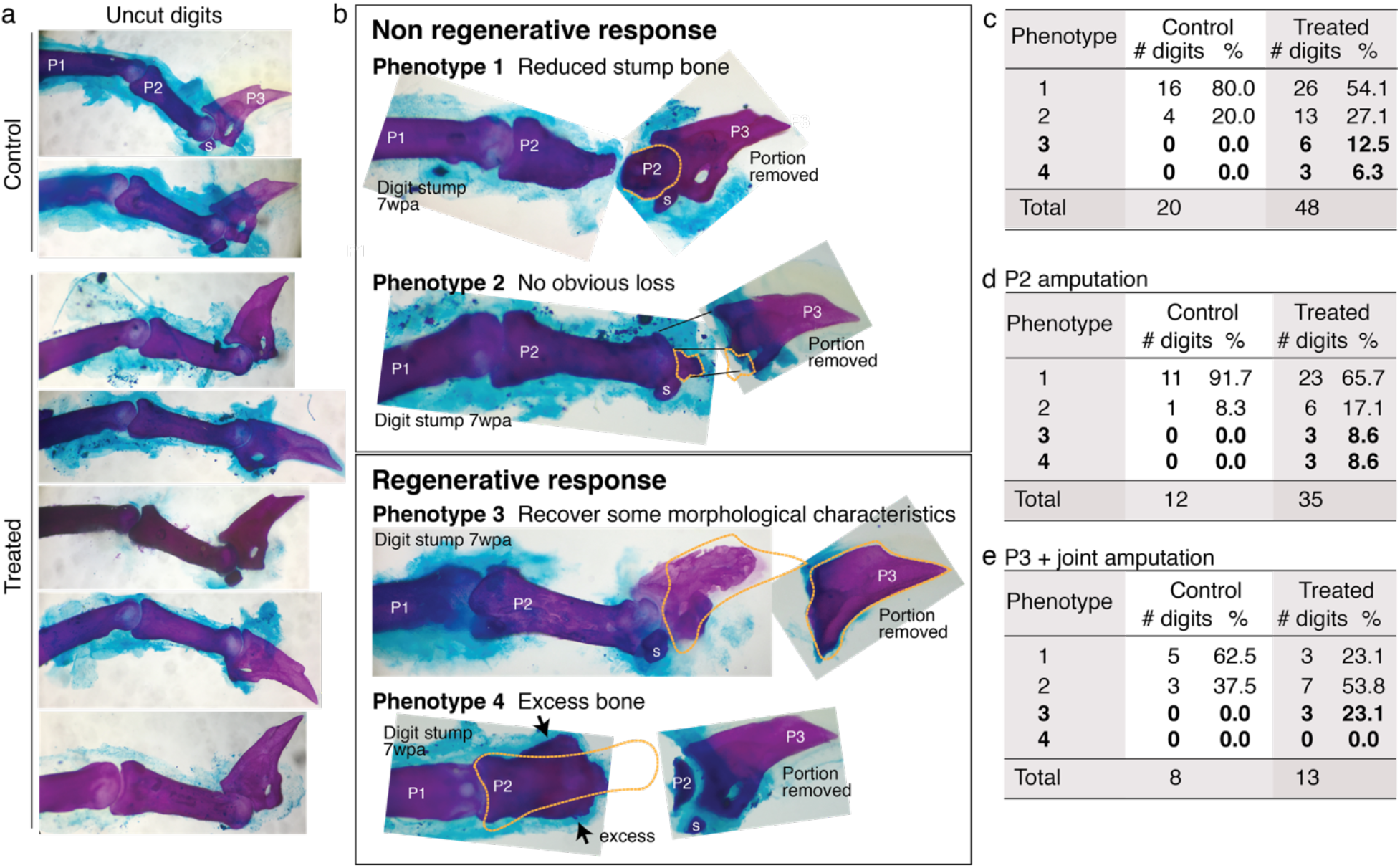
Mouse digit phenotypes. Whole-mount skeletal staining was performed with Alizarin red. wpa: week post amputation, P1: phalange 1, P2: phalange 2, P3: phalange 3, s: sesamoid bone **(a)** Skeletal staining of unamputated digits (digit 3) from control and treated groups show no obvious differences in uncut digits due to the treatment. **(b)** Skeletal staining of digits stumps at 7 wpa and the original portion removed from the digits. Some digit stumps show no change or appear to have undergone histolysis (Chamberlain et al. 2017) resulting in reduced bone mass (Phenotype 1 and 2). Some digit stumps show regenerative response, either recovery of some morphological characteristics (Phenotype 3, detailed more in Figure 6**—**figure supplement 2) or excess, ectopic bone mass (Phenotype 4). We erred on the conservative side in scoring phenotype 3 and 4; when in doubt, digits were classified into phenotype 1 or 2. **(c-e)** Phenotype counts in all digits **(c)**, in digits amputated across P2 **(d)**, and in digits amputated across P3 or joint **(e)**.

**Figure S11.**
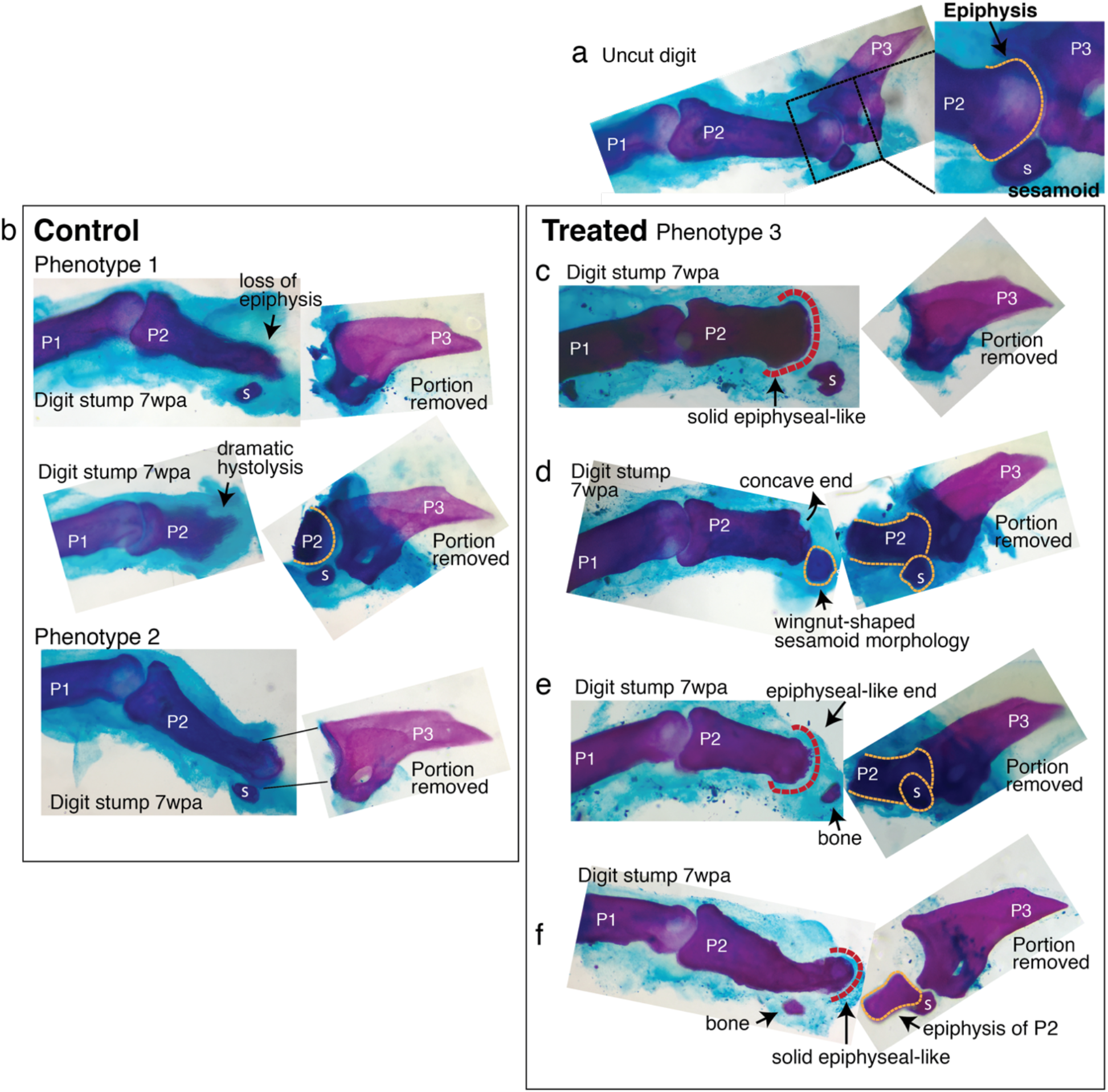
Regenerative response observed in mouse digit. Six digit stumps (of total 48 examined) show regenerative response. The most dramatic two are presented in Figure 4. The remaining four are presented here. wpa: week post amputation, P1: phalange 1, P2: phalange 2, P3: phalange 3, s: sesamoid bone **(a)** An uncut digit, shown for a comparison. Magnified is the P2/P3 joint area to highlight key morphological markers: the knobby epiphyseal cap of P2 and the sesamoid bone embedded in the tendon on the flexor side of P2. **(b)** Digit stumps from control mice show either bone stump histolysis (top and middle, phenotype 1) and no visible changes in bone stump (bottom, phenotype 2). **(c-f)** Digit stumps from treated mice that show regenerative response. **(c)** In this digit, the amputation removed all P3 by a cut through the joint. At 7 wpa, the P2 stump is reduced, but recovered the epiphyseal-like end (red dashed line) — marked by solid curved shape, as opposed to irregularly shaped histolyzing bone. **(d)** In this digit, the amputation removed a significant portion of P2 and the sesamoid bone. The P2 stump does not regain an epiphyseal end (the end is concave and irregular). However, the sesamoid bone is reformed, as identified by its location on the flexor side of P2 and wingnut shape (Wirtschafter and Tsujimura, 1961) under the microscope. The recovery of sesamoid bone is non-trivial, as digit sesamoids form in juxtaposition to the condensing phalange, detaching from the phalange by formation of a cartilaginous joint (Eyal et al. 2019). **(e)** In this digit, the amputation removed a significant portion of P2 and the sesamoid bone. At 7 wpa, the P2 stump appears to be reforming an epiphyseal, rounded end (red dashed line). There is a small bone distal to P2, whose curvature articulates with the P2 end, but there are not enough morphological characters to identify the bone. **(f)** In this digit, the amputation removed the epiphyseal cap of P2 and the sesamoid bone. The P2 stump appears to have lost some mass, but reforms an epiphyseal-like end (red dashed line). There is an additional small bone located where the sesamoid bone should be, but lacks sufficient morphological characters to identify.

## Methods

### Aurelia aurita

The experiments were performed in *Aurelia aurita* sp. 1 strain, also alternatively named *Aurelia coerulea* based on recent molecular classification (Scorrano et al., 2016). Polyps were reared at 68°F, in 32 ppt artificial seawater (ASW, Instant Ocean), and fed daily with brine shrimps (*Artemia nauplii*) enriched with *Nannochloropsis* algae (both from Brine Shrimp Direct). To induce strobilation, polyps were incubated in 25 M 5-methoxy-2-methyl-indole (Sigma M15451) at 68°F for an hour (Fuchs et. al, 2014).40 Ephyrae typically began to strobilate within a week.

### Amputation

Strobilated ephyrae were fed daily with rotifers (*Brachionus plicatilis*, Reed Mariculture) until amputation time. 2-3 days old ephyrae were anesthetized in 400 μM menthol and amputated using a razor blade mounted on an x-acto knife handle. After amputation, ephyrae were let to recover in bubbler cones (**Figure S4**). Regeneration was assessed at various times for 1-2 weeks after amputation, before onset of maturation to medusa.

### Experiment in the original habitat

The polyp population in the study arose from parental polyps collected off the coast of Long Beach, CA (33°46’04.2”N 118°07’44.2”W,_GPS: 33.7678376,-118.1289559). Ephyrae were amputated in location and immediately after submersed in the ocean. For submerging the amputated ephyrae in the ocean, a two-layered aquarium was custom-built. Ephyrae were placed in plastic canisters with a 7 cm diameter hole cut in the lid and covered with a 250 μm plastic screen. The canisters were then placed in a thick plastic tank fitted with a 500 μm plastic screen on top. This design offers protection to the ephyrae against predators and strong waves, while at the same time allowing exchange of water, zooplanktons, and other particulates. Ephyrae were collected after two weeks.

### Regeneration experiments

All experiments were performed at 68°F. Amputated ephyrae were let to recover in 1 L sand settling cones (Nalgene Imhoff, **Figure S4**). In each cone, an airline from a Tetra Whisper 100 pump was placed at the bottom to create a gentle upward current (~1 air bubble/sec, **Movie S2**). In this “bubbler cone” setup, the ephyrae continually experienced water current, either the upward bubble-generated current or the downward gravity-generated current. The conical geometry helps avoid stagnant spots, where the ephyrae could get stuck. Each cone housed 30 ephyrae in 500 mL ASW to avoid crowding and fouling. ASW was changed weekly.

### Nutrients

Amputated ephyrae were fed daily with rotifers. The number of rotifers was estimated using a 6-well plate fitted with STEMgrid™ (the same principle as using a hemocytometer). In this study, low food was ~100-200 rotifers/ephyra and high food was 400 rotifers/ephyra. To replicate the study, these numbers should only be used as initial estimates, as what is “low” or “high” food amount may be relative to and easily vary across lab cultures (*e.g.*, rotifer culture, differences across *Aurelia* strains, *etc.*). Most if not all rotifers were typically consumed within an hour (determined by measuring the rotifers in the water).

### Insulin

Immediately after amputation, ephyrae were placed in ASW supplemented with 500 nM human recombinant insulin (Sigma I0908). Insulin was refreshed weekly. To determine the concentration used, a range of concentrations, 10 nM to 3 mM, were tested. The concentration 500 nM was chosen as it maximized regeneration frequency while avoiding solubility problems. To control that the effect of insulin was not due to non-specific additions of proteins, BSA at 500 nM and 3 mM were tested.

### Hypoxia

Immediately after amputation, ephyrae were placed in hypoxic ASW. To create a hypoxic environment, nitrogen or argon, instead of ambient air, was pumped into the bubbler cone, beginning from the day before the experiment and maintained throughout the duration of the experiment. The bubbler cone was sealed with parafilm to maintain the lowered oxygen level. The nitrogen/argon flow was adjusted to achieve 50% reduction in the dissolved oxygen level. Dissolved oxygen level was measured using a Clark-type electrode Unisense OX-500 microsensor. The measurement was normalized to oxygen level in control ASW bubbled normally with ambient air. Oxygen measurement was performed prior to the experiment and subsequently every 3 days.

### L-leucine

Immediately after amputation, ephyrae were placed in ASW supplemented with 100 M L-leucine (Sigma L1002, the cell-permeable methyl ester hydrochloride form). L-leucine was refreshed weekly. To determine the concentration used, a range of concentrations from one to hundreds of mM was tested. The concentration of 100 mM was chosen as it maximized the regeneration frequency without non-specific, negative effects.

### Statistical analysis

To assess the statistical significance of the treatments, meta-analysis of effect size was performed (Borenstein et al., 2009). For each experiment, the effect size of a treatment was computed relative to the internal control set up using ephyrae from the same clutch. The effect size metrics used are determined by the form of the dataset. For measurements of frequencies (*e.g.*, regeneration frequency), the datasets are in the form of a 2 × 2 table of dichotomous variables,

|  | # ephyrae that regenerate | # ephyrae that do not regenerate |
| --- | --- | --- |
| Control | a | b |
| Treatment | c | d |

For such 2 × 2 datasets, in situations where the baseline varies (*e.g.*, varying baseline regeneration across clutches), the commonly used measures of effect size are the Risk Ratio (RR),

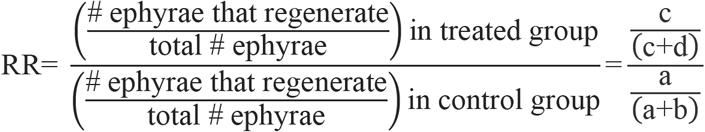

and the Odds Ratio (OR),

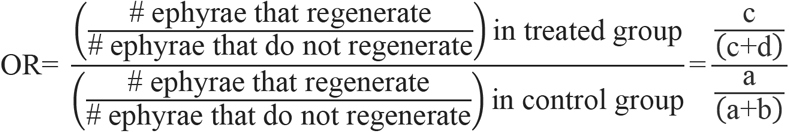

RR compares the probability of an outcome in treated vs control group, whereas OR compares the odds of an outcome in treated vs control group.

For measurements of arm length and body size, the datasets are in the form of continuous variables. For such data, the commonly used effect size is the Response Ratio (R),

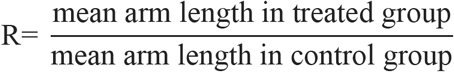

R evaluates the proportionate change that results from a treatment, and is the meaningful effect size to use when the outcome of a treatment is measured on a physical scale, *e.g.*, length or area (as opposed to arbitrary scale, *e.g.*, happiness level). Experiments where regeneration in one of the groups occurred in 0 ephyra were necessarily excluded.

Having computed the effect size (RR, OR, or R) within each experiment, meta-analysis of the effect size across experiments was performed. The metafor package^15^ in R was used, with fixed-effect model (for nutrients and leucine) or random-effect restricted maximum likelihood model (for insulin and hypoxia, which had different control conditions across the experiments). Statistical coefficients were based on normal distribution.

### Phalloidin and tyrosinated tubulin staining

All steps were performed at room temperature, unless indicated otherwise. Ephyrae were first anesthetized in 400 μM menthol, which minimizes curling during fixing. Next, ephyrae were fixed in 3.7% (v/v) formaldehyde (in PBS) for 15 minutes, permeabilized in 0.5% Triton X-100 (in PBS) for 5 minutes, and blocked in 3% (w/v) BSA for 2 minutes. For neuron staining, ephyrae were incubated in 1:200 mouse anti-tyrosinated alpha tubulin antibody (Sigma MAB1864-I) overnight at 4°C, and then in 1:200 goat-anti-mouse Alexa Fluor 488 (Sigma A11029) overnight in the dark at 4°C. Primary or secondary antibodies were diluted in 3% BSA. For actin staining, ephyrae were incubated in 1:20 Alexa Fluor 555 Phalloidin (Life Technologies A12379) overnight or for 2 hours in the dark at 4°C. For nuclei staining, ephyrae were incubated in 1:10 Hoechst 33342 (Sigma B2261) for 30 minutes in the dark.

**Microscopy._** Ephyrae were imaged anesthetized in menthol._Brightfield images, fluorescent images, and movies were taken with the Zeiss AxioZoom.V16 stereo zoom microscope and AxioCam HR 13-megapixel camera. Optical sectioning was performed with ApoTome.2.

### Drosophila melanogaster

OregonR and CantonS wild type strains were reared under standard conditions at 23°C.

### Amputation

Amputation was performed on adult flies 2-7 days after eclosion. Flies were anesthetized with CO_2_, placed under a dissection microscope, and tibia amputated using a spring scissors (Fine Science Tools, 91500-09) and superfine dissecting forceps (VWR, 82027-402). See **Figure 4** for detailed description of the amputation plane. Recovering *Drosophila* were fed with standard lab fly food (control) or standard lab fly food mixed with 5 mM L-Leucine (Sigma L8000), 5 mM L-Glutamine (Sigma G3126), and 0.1 mg/mL insulin (human recombinant, MP Biomedicals 0219390080). To introduce the molecular factors, the fly food was microwaved in short pulses, such that the topmost layer of the food was liquified. Molecular factors in aqueous medium were then pipetted into this liquified layer. Food was allowed to re-set at 4°C for at least 20 minutes. New food was prepared fresh every 2 days, and flies were moved into freshly prepared treated food every 2 days, throughout the course of the 2- to 3-week experiment. The *Drosophila* data reported in this study were reproduced by 3 independent experimenters, with many experiments examined at multiple times by 2 experimenters.

### DAPI staining

Fly tibias were dissected and washed in 70% ethanol (<1min) to decrease the hydrophobicity of the cuticle and washed in PBS with 0.3% Triton-X for 10 minutes. The legs were fixed in 4% paraformaldehyde (in PBS) overnight at 4°C and washed five times for 20 minutes each in PBS with 0.3% Triton-X. The legs were equilibrated in Vectashield mounting medium with DAPI (Vector H-1200) overnight at 4°C, and imaged using Zeiss AxioZoom.V16 stereo zoom microscope with AxioCam HR 13-megapixel camera. Confocal imaging was performed using X-Light V2 spinning disk mounted on the Olympus IX81 inverted microscope.

### Live fly imaging

Flies anesthetized on a CO2 bed were imaged under a dissection scope equipped with the Zeiss AxioCam 503 color camera.

### Electron microscopy

Environmental scanning electron microscopy (ESEM) was performed on a FEI Quanta 200F (FEI, Hillsboro, Oregon). Whole live flies were mounted onto the SEM stub with copper tape. ESEM images were attained at a pressure of 0.1 mbar and 5 kV at a working distance of 9-12 mm, with water as the ionizing gas.

### Mus musculus

All studies comply with relevant ethical regulations for animal testing and research, and received ethical approval by the Institutional Animal Care and Use Committees at the California Institute of Technology.

### Strain

Adult female (3-6 months old) wild-type CD1 mice (Charles River Laboratories strain 022) were used for all regeneration studies.

### Mouse digit amputation

Digit amputation was performed following the established protocol in the field (Simkin et al., 2013). Mice were anesthetized with 1-5% isoflurane (in oxygen) in an induction chamber, followed by maintenance on a nosecone. The mouse was positioned on its belly with its hind paws outstretched and the ventral side of the paw facing upwards. Sustained-Release Buprenorphine was administered (Buprenorphine SR LAB®) at 0.5 mg/kg subcutaneously as an analgesic. Blood flow to the hindlimb was stemmed by tying a rubber band around the ankle and clamping it with a hemostat. All surgical procedures were carried out under a Zeiss Stemi 305 dissection microscope. An initial incision, parallel to the position of foot, was made through the ventral fat pad using Vannas spring scissors (World Precision Instruments, 14003). The length of this incision was determined by the amount of ventral skin needed to seal the digit amputation wound completely. The ventral skin freed in the initial incision was peeled back using surgical forceps, and a no. 10 scalpel (Sklar, 06-3110) was used to amputate and bisect the digit completely through the second or third phalange. Digits 2 and 4 on the right hind paw were operated on in this fashion, while digit 3 remained unamputated as a control. The amputation wound was immediately closed with the ventral skin flap and sealed with GLUture (Zoetis, Kalamazoo, MI). Amputated portions were immediately fixed as control for skeletal staining. Amputated digits were photographed weekly for 7 weeks, at which time the digits were dissected for skeletal staining.

### Mouse digit dissection and skeletal staining

Mice were euthanized and digits 2, 3 and 4 were removed with a no. 10 scalpel (Sklar, 06-3110) through the first phalange. Excess skin and flesh were removed with spring scissors (Fine Science Tools, 91500-09) and fine dissecting forceps (Fine Science Tools, 11254-20). All digits analyzed by whole-mount skeletal stains were prepared with a standard alizarin red and alcian blue staining protocol.48 Digits were dehydrated in 95% ethanol for 1 day, and incubated in staining solution (0.005% alizarin red, 0.015% alcian blue, 5% acetic acid, 60% ethanol) for 1 day at 37°C. Tissue was cleared in 2% potassium hydroxide at room temperature for 1 day, 1% potassium hydroxide for 1 day, and then taken through an increasing glycerol series (25%, 50%, 75%, 100%). The stained samples were imaged on Zeiss AxioZoom.V16 stereo zoom microscope with a Zeiss AxioCam 503 color camera or a Zeiss Stemi 305 dissection microscope with an iPhone 6 camera.

### Data Availability

Raw image raw data from the regeneration induction experiments (*i.e.*, images of ephyrae from the main experiments in Figure 4, and all mouse digits analyzed) are deposited in the public repository Image Data Resource (http://idr.openmicroscopy.org/about/). Supporting raw data are available upon request.

